# COUP-TFI/Nr2f1 orchestrates intrinsic neuronal activity during cortical area patterning

**DOI:** 10.1101/728402

**Authors:** Isabel del Pino, Chiara Tocco, Elia Magrinelli, Andrea Marcantoni, Celeste Ferraguto, Giulia Tomagra, Michele Bertacchi, Christian Alfano, Xavier Leinekugel, Andreas Frick, Michèle Studer

## Abstract

The formation of functional cortical maps in the cerebral cortex results from a timely regulated interaction between intrinsic genetic mechanisms and electrical activity. To understand how transcriptional regulation influences network activity and neuronal excitability within the neocortex, we used mice deficient for the area mapping gene *Nr2f1* (also known as *COUP-TFI*), a key determinant of somatosensory area specification during development. We found that cortical loss of *Nr2f1* impacts on spontaneous network activity and synchronization at perinatal stages. In addition, we observed alterations in the intrinsic excitability and morphological features of layer V pyramidal neurons. Accordingly, we identified distinct voltage-gated ion channels regulated by *Nr2f1* that might directly influence intrinsic bioelectrical properties during critical time windows of somatosensory cortex specification. Together, our data suggest a tight link between *Nr2f1* and neuronal excitability in the developmental sequence that ultimately sculpts the emergence of cortical network activity within the immature neocortex.

## INTRODUCTION

The area- and cell-type specific organization of the mammalian neocortex is a process involving coordinated interactions between cell intrinsic genetic programs and neural activity, an essential feature in specifying the composition and organization of neural circuits during all stages of development (Jabaudon, 2017; Simi and Studer, 2018). Before experience shapes neuronal circuits via sensory-driven activity, the developing cortex is genetically primed to establish patterns of local spontaneous activity, which autonomously synchronizes large groups of neurons independently of sensory stimulation and contributes to the development of neuronal circuits (Luhmann and Khazipov, 2018; Andreae and Burrone, 2018; Kirischuk *et al*., 2017; Anton-Bolanos *et al*., 2019). These early patterns of spontaneous activity are thought to set the initial blueprint of cortical network organization, and their alteration during cortical development leads to the formation of dysfunctional cortical circuits (Kirkby *et al*., 2013; Li *et al*., 2013). The embryonic thalamus also conveys spontaneous neuronal activity to the immature cortex, influencing the formation of a functional somatotopic map (Anton-Bolanos *et al*., 2019). Hence, local spontaneous activity in cortical and pre-cortical relay stations during embryonic and early postnatal stages might prepare cortical areas and circuits for upcoming sensory input.

Spontaneous activity emerging during early development in different cortical areas in form of calcium (Ca^2+^) transients have been reported already at embryonic day E16 in mice, most probably generated in proliferating radial glia cells, as slowly propagating calcium waves (Kirischuk *et al*., 2017). Immature neurons express voltage-dependent ion channels, responsible for a highly synchronized spontaneous activity within small neuronal networks (Luhmann *et al*., 2016). Finally, certain features of immature cells, including high membrane resistance and prominent low-threshold calcium currents, seem to enhance excitability in neurons receiving little afferent sensory input (Moody and Bosma, 2005).

Besides spontaneous activity, the neocortical primordium is also under the control of distinct morphogens that establish proper coordinates along the developing anteroposterior and dorsoventral cortical axes (Greig *et al*., 2013; O’Leary and Sahara, 2008; Alfano and Studer, 2012). Functioning in a dose-, context- and time-dependent manner, these signaling pathways modulate gradient expression of area patterning genes, such as *Pax6*, *Sp8*, *Emx2* and *Nr2f1* (also known as COUP-TFI), among others. These transcriptional regulators cooperatively regulate cell identity specification in cortical progenitors and early postmitotic neurons, building a rough primordial areal map, known as a protomap, which will be refined by sensory-evoked activity during postnatal stages (Simi and Studer, 2018). Alterations in genetic dosage of these factors alter the size and position of primordial area maps with severe consequences in cell-type specification and circuit formation (Greig *et al*., 2013; Jabaudon, 2017). Although the interaction between cortical patterning genes and onset of spontaneous activity during primary area mapping has been hypothesized (Simi and Studer, 2018), there is still no clear evidence on how this interaction is perpetrated.

To tackle this question, we used mice deficient for the nuclear receptor *Nr2f1*, a key transcriptional regulator during cortical development, as a genetic model of altered somatosensory (S1) area mapping (Armentano *et al*., 2007). As a member of the superfamily of steroid hormone receptors, *Nr2f1* represents a well-studied area patterning gene, which determines sensory identity predominantly in early postmitotic cells during neocortical area mapping, and controls neuronal migration and differentiation (Alfano *et al*., 2014; Tomassy *et al*., 2010; Alfano *et al*., 2011; Faedo *et al*., 2008). Haploinsufficiency of the human *NR2F1* gene leads to an emerging genetic neurodevelopmental disorder, named Bosch-Boonstra-Schaaf Optic Atrophy Syndrome (BBSOAS). Patients affected by this autosomal-dominant disorder exhibit several clinical features, such as intellectual disability, epileptic seizures, autistic-like traits, abnormal fine motor coordination and optic atrophy, among others (Bosch *et al*., 2014; Chen *et al*., 2016). In mice, *Nr2f1* deficiency solely in the cortex prevents the formation of a somatotopic map, affects radial migration and laminar specification, ultimately leading to distinct motor and cognitive impairments (Alfano *et al*., 2014; Armentano *et al*., 2007; Tomassy *et al*., 2010; Contesse *et al*., 2019). Thalamocortical axons reach the presumptive S1 cortex at prenatal stages in *Nr2f1* mutants, but fail to functionally innervate the cortical plate at postnatal stages (Armentano *et al*., 2007), suggesting that loss of Nr2f1 function within the cortex prevents the arrival of sensory peripheral inputs in these circuits. In this study, we show that cortices deficient for Nr2f1 have altered Ca^2+^ burst frequency and network firing activity at perinatal stages, before sensory evoked activity has taken place. We also observed that glutamatergic pyramidal neurons, particularly those of layer V, display changes in intrinsic membrane excitability along with neuroanatomical and structural alterations within the immature cortex. Finally, we found that altered levels of several voltage-gated ion channels including HCN1 might underlie an aberrant state of neural excitability in the early postnatal neocortex. Together, our data show a novel role for Nr2f1 in controlling cortical area patterning through the modulation of spontaneous activity intrinsic to the neocortex. This newly described function is accomplished by the transcriptional modulation of a plethora of channels involved in modulating intrinsic excitability as well as morphological and structural features of layer V pyramidal neurons.

## MATERIALS AND METHODS

### Mice

*Nr2f1*/COUP-TFI*fl/fl* mice were crossed to *Emx1-Cre* mice to delete *COUP-TFI/Nr2f1* exclusively in cortical progenitors and their progeny (Armentano *et al*., 2007). These mice are named *Nr2f1cKO* throughout the text. Mice were genotyped as previously described (Armentano *et al*., 2007). Control and mutant littermates were *COUP-TFIfl/fl* and *COUP-TFIfl/fl:Emx1-Cre*, respectively. For P21 studies, control (*Nr2f1 fl/fl)* and mutant *Nr2f1cKO* mice were crossed to *Thy1-eYFP-H* mice to specifically mark LVPNs, as previously reported (Harb *et al*., 2016). Midday of the day of the vaginal plug was considered as embryonic day 0.5 (E0.5). Control (*Nr2f1 fl/fl)* and mutant *Nr2f1cKO* mice were predominantly bred in a C57BL6 background. Both male and female embryos and pups were used in this study. All mouse experiments were conducted according to national and international guidelines, with authorization #15349 and 15350 by the French Ministry of Education, Research and Innovation (MESRI) and upon request of the local ethics committee – CIEPAL NCE/2019-548 (Nice) and CEEA50 (Bordeaux), and in accordance with the guidelines established by the Italian National Council on Animal Care and approved by the local Animal Care Committee of University of Turin (Authorization DGSAF 0011710-P-26/07/2017).

### MEA recordings on primary cultures and analysis of MEA activity

Cortical neurons were obtained from E18.5 *Ctrl* and *Nr2f1cKO* mouse embryos. Somatosensory cortices were rapidly dissected under sterile conditions, kept in cold HBSS (4°C) with high glucose, and then digested with papain (0.5 mg/ml) dissolved in HBSS plus DNAse (0.1 mg/ml), as previously described (Gavello *et al*., 2018). Isolated cells were then plated at the final density of 1200 cells/mm^2^ onto the MEA (previously coated with poly-DL-lysine and laminin). Cells were incubated with 1 % penicillin/streptomycin, 1% glutamax, 2.5% foetal bovine serum, 2% B-27 supplemented neurobasal medium in a humidified 5 % CO_2_ atmosphere at 37°C. Each MEA dish was covered with a fluorinated ethylene-propylene membrane (ALA scientific, Westbury, NY, USA) to reduce medium evaporation and maintain sterility, thus allowing repeated recordings from the same chip.

Multisite extracellular recordings from 60 electrodes were performed using the MEA-system, purchased from Multi-Channel Systems (Reutlingen, Germany). Data acquisition was controlled through MC_Rack software (Multi-Channel Systems Reutlingen, Germany), setting the threshold for spike detection at -30 µV and sampling at 10 kHz. Experiments were performed in a non-humidified incubator at 37°C and with 5% CO_2,_ without replacing the culture medium. Before starting the experiments, cells were allowed to stabilize in the non-humified incubator for 90 seconds; then the spontaneous activity was recorded for 2 minutes.

Bursts analysis was performed using Neuroexplorer software (Nex Technologies, Littleton, MA, USA) after spike sorting operations. A burst consists of a group of spikes with decreasing amplitude, thus we set a threshold of at least 3 spikes and a minimum burst duration of 10ms. We set interval algorithm specifications such as maximum interval to start burst (0.17 sec) and maximum interval to end burst (0.3 sec) (Gavello *et al*., 2012; Gavello *et al*., 2018). Burst analysis was performed by monitoring the following parameters: mean frequency, number of bursts and mean burst duration. Cross correlation probability values were obtained by means of Neuroexplorer software using ± 2 s and 5 ms bin size. The number of experiments is given considering the number of MEA (N_MEA_), each of those consisting of 59 recording electrodes. Where indicated we provided the total number of recording electrodes (N_channels_).

### Ca^2+^ imaging from *ex vivo* brain slices preparation and analysis

Animals were anesthetized by cold and brains were dissected in ice cold high-Mg artificial cerebral-spinal fluid (aCSF; 125mM NaCl, 25mM NaHCO_3_, 2.5mM KCl, 7mM MgCl_2_, 2mM CaCl_2_, 1.25mM NaH_2_PO_4_, 10mM glucose, pH7.4) after decapitation. Horizontal 300 µm slices were prepared on a Leica Vibratome 100M in ice cold High Mg ACSF under 95% O_2_, 5% CO_2_ bubbling. Selected slices where incubated 1h at 37°C in ACSF (125mM NaCl, 25mM NaHCO_3_, 2.5mM KCl, 2mM CaCl_2_, 1.25mM NaH_2_PO_4_, 10mM glucose, pH7.4) 0.63% Oregon Green BAPTA^TM^ (ThermoFisher O6807) in DMSO before proceeding to image acquisition. Slices were loaded on a 35mm diameter petri dish with 1.3mm glass bottom and immobilized with a custom nylon net while kept under constant aCSF perfusion at 37°C throughout the imaging process. After 30 minutes of equilibration in the microscope objective holder time lapse images were acquired over a Spinning Disk microscope (Olympus / Andor technology) with a 40x air objective, recording for 15 minutes at 150 ms intervals.

Images were automatically corrected for drifting using the built-in Fiji function “*Linear stack alignment with SIFT*” (Schindelin *et al*., 2012). Single cell borders were manually generated as separate ROI (Regions of Interest) on a custom MATLAB script. Each ROI was used to acquire BAPTA fluorescence intensity as the mean of intensity within each ROI area by frame, resulting in a matrix of single cell time series of BAPTA fluorescence signals as a readout of Ca^2+^ cytoplasmic concentration. Matrices were corrected by moving average (Anderson, 1975) normalizing each timepoint with the average of 60 seconds for each cell to generate a constant baseline over the progressive signal loss due to the progressive BAPTA bleaching. As a cut off to detect actively bursting cells, we discarded from the analysis cells whose distribution of intensity for each time frame arranged more than 40% of the frames above the first quarter of maximum corrected fluorescence intensity. For the remaining selected timeseries, Ca^2+^bursts were detected filtering BAPTA intensity on a threshold of 3 standard deviations. The bursting frequency of every selected cell was measured dividing the number of bursts above threshold by the sum of intervals between each burst. To evaluate coordination of activity, we counted the ratio of bursts occurring across cells in an interval of 3s normalized by the total number of bursts. All time-lapse analysis was performed using a custom MATLAB script.

### *Ex vivo* electrophysiology and analysis

Mouse pups at postnatal stage (P)5–P8 were decapitated and brains were placed on ice-cold modified artificial cerebrospinal fluid (aCSF) containing (in mM): 87 NaCl, 25 NaHCO_3_, 5 Glucose, 65 Sucrose, 2.5 KCl, 1.25 NaH_2_PO_4_, 0.5 CaCl_2_, 7 MgCl_2_ saturated with 95% CO_2_ and 5% O_2_. Coronal slices of 300µm were cut in a vibratome (Vibratome 300 Plus, Sectioning Systems) and gently transferred to room temperature aCSF containing (in mM): 125 NaCl, 25 NaHCO_3_, 25 Glucose, 2.5 KCl, 1.25 NaH_2_PO_4_, 2 CaCl_2_, 1 MgCl_2_ saturated with 95% CO_2_ and 5% O_2_ (pH 7.4). Current-clamp recordings were performed at 32-34°C using a potassium-gluconate based intracellular solution containing (in mM): 120 K-Gluconate, 20 HEPES, 0.5 EGTA, 15 KCl, 4 MgATP and 0.3 NaGTP (pH 7.4) and osmolarity set to 299 mosmol l^-1^. Biocytin (1.5 – 2 mg ml^-1^) was added to the intracellular solution for post-hoc immunohistochemistry. Neurons were visualized with an upright microscope (Zeiss Axio Examiner D1) equipped with an Evolve 512 EMCCD Camera (Photometrics) using a 63X/1.0nA water-immersion objective (Zeiss) and infrared DodT contrast imaging. Recording pipettes (6-10MΩ) were pulled from borosilicate glass using a Narishige P-10 puller. Data was acquired using a Dagan BVC-700A Amplifier, low-pass filtered at 3kHz and sampled at 20kHz with ITC16 (Instrutech) and AxoGraph X software. Membrane voltage was recorded in the bridge mode. To determine the excitability of pyramidal neurons, we performed 800ms square current steps of 10pA from -100pA to +150pA. The rheobase was defined as the minimum somatic current required to elicit an action potential (AP) from the resting membrane potential of the neuron. Input resistance was calculated from voltage responses to -20pA. Threshold potential was defined as *dV/dt* = 10mV/ms. Inter-spike interval (ISI) was calculated as the difference between first and second AP at the minimal current to elicit >2AP. The amplitude of action potentials was measured from spike threshold (see Table 1). Action potential width was measured at half of the maximum amplitude (APhalf-width or AP*h-w*). Voltage sag was calculated as the difference between negative voltage peak and steady-state hyperpolarization from traces in which the difference between steady-state response and baseline fell within 20-30mV.

### *Post-hoc* immunofluorescence

Brain slices were fixed overnight at 4°C with 4% paraformaldehyde in 0.1M phosphate buffer (PB) pH7.4. The following day slices were washed 3 x 10 min with 0.1M PB and incubated overnight at 4°C with Alexa-Fluor®-conjugated streptavidin diluted 1:500 in 0.1M PB. The following day slices were washed 3 x 10 min with 0.1M PB, stained with DAPI and mounted with Mowiol.

### KCl *ex vivo* depolarization assay

*Ctrl* and *Nr2f1cKO* P7 mice were sacrificed by decapitation and brains dissected into ice-cold aCSF saturated with 95% O_2_ and 5% CO_2_. 300µm-thick slices were produced by vibratome sectioning and then transferred into homemade floating nests immersed into aCSF, kept at room temperature and constantly saturated with 95% O_2_ and 5% CO_2_. After 15 minutes of acclimatization, slices from both genotypes were split into two groups: an untreated group (**-KCl**) and an experimental group (**+ KCl**). KCl were added to the aCSF of the experimental group whereas control slices were kept in normal aCSF. After an incubation time of 6 hours(Dumitrescu, Evans and Grubb, 2016), slices were washed for 15 minutes in aCSF and then processed for immunofluorescence (IF), as described below.

### Histology and immunohistochemistry of mouse perinatal cortices

Perinatal mice (P7-8) were anesthetized by using a mixture of Tiletamine-Zolazepam-Xylazine-Buprenorphine (TZXB) and intracardially perfused with Phosphate Buffer Saline (PBS) followed by 4% paraformaldehyde (PFA) in PBS. Then, brains were dissected and post-fixed for 4h at 4°C in 4% PFA and then vibratome-sectioned to obtain 100 m thick, coronal or sagittal floating sections. Overnight (ON) incubation at 4°C in a solution of 0.5% Triton X-100, 3% BSA, 10% Goat Serum in PBS was performed to permeabilize the sections and reduce non-specific binding of the antibodies. For IF, sections were incubated for two days at 4°C with indicated primary antibodies in a solution of 0.5% Triton X-100, 10% goat serum in PBS and then ON at 4°C with relative secondary antibodies (see **Table S3**) and HOECHST diluted in PBS. For HCN1 IF, brains were briefly post-fixed for 30 minutes (min) at 4°C in 4% PFA and then immediately sectioned with the vibratome to obtain 50µm thick coronal floating sections. Permeabilization-blocking step was reduced to 2h, primary antibody incubation to one ON and secondary antibody and HOECHST incubation to 2h. Sections were then transferred on superfrost plus slides (ThermoScientific), let dry for 30min to 1h and finally mounted with Mowiol mounting medium.

### Microscopy and image analysis

Imaging was performed using a Zeiss 710 confocal microscope equipped with a 405nm diode, an argon ion, a 561nm DPSS and a 647 HeNe lasers. Z-stacks of fixed cortical sections were imaged using a LD-LCI Plan-Apo 25x/0.8 NA or a LD Plan-Apo 63x/1.4 NA oil immersion lenses. Images from IF experiments were analyzed by using Fiji-ImageJ Software (Schindelin *et al*., 2012): cell counter plug-in was used to assess the number of cells expressing specific proteins, while measurement tools were used to determine signal intensity or Axonal Initial Segment (AIS) features such as length and distance from the soma. For the AIS average diameter measurement, we took advantage of IMARIS software’s tool filament tracer. AISs were semi-automatically reconstructed and an average diameter automatically measured for every AIS individually.

Morphological characterization of biocytin filled layer V pyramidal neurons (LVPNs) was analyzed by using IMARIS software. Cell arborization was reconstructed in a semi-automatic manner, via the filament tracer module. Automatic length measuring tool was used to calculate the total dendritic length and the basal dendritic length. Branching points, branching point orders and tips were manually quantified for both basal and apical dendrites, as well as primary and lateral dendrites. The branch tip order is an integer value equivalent to the number of branching points a dendrite undergoes from its somatic origin to the terminal tip (see also **Figure 4D** for a graphic representation). Primary dendrites were identified as directly originating from the soma, lateral dendrites were categorized as originating from and perpendicular to the apical dendrite whereas total and basal branch tips were identified as the terminal ends of primary dendrites including or excluding the apical dendrite arborization respectively. The basal Dendritic Complexity Index (bDCI) was calculated as follow:

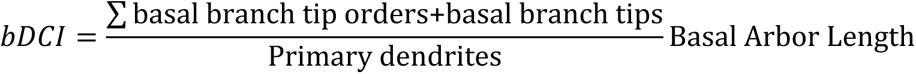

An automatic detection of Sholl intersections was used to compute Sholl profiles for every individual cells. Sholl intersections were identified as the number of dendrite intersections for concentric spheres of increasing radius (10µm difference) and having as a center the centroid of the cell body.

### RNA extraction and cDNA synthesis

RNA for reverse transcriptase–PCRs (RT–PCRs) was extracted from P7 dissected neocortices separated from meninges of *Ctrl* and *Nr2f1cKO* mice (n=3) using the RNeasy Mini kit (Qiagen) following manufacturer’s instructions. We synthetized cDNA from 1μg of total RNA using Superscript III First-Strand Synthesis System for RT–PCR (Invitrogen) following manufacturer’s instructions.

### qPCR RNA/cDNA quantification

qPCRs on cDNA was performed using KAPA Sybr Fast (Kapa Biosystems) according to manufacturer’s instructions. Samples were run on a LigthCycler II 480 (Roche) and the following primers were used: GAPDH Fw: GTATGACTGCACTCACGGCAAA. Rev: TTCCCATTCTCGGCCTTG; Cacna1A Fw: GGGCCCAAGAGATGTTCCAG. Rev: TCCACAGACTGGGAGTTGGG; Cacna1B Fw: CACGCAGAGGACCCACGATG. Rev: CATCACAGCCAGTGTTCCTG; Cacna1G Fw: TCAGCTGCCTGTCAACTCCC. Rev: CCCATCACCATCCACACTGG; Cacna1H Fw: ACCTACACAGGCCCGGTCAC. Rev: ATGGGACCTGGAAGGAGGTG; Cacna2D1 Fw: CAGGGAGGGCACTGATCTTC. Rev: TTGATAGTGACGGGCGAAGG; Cacna2D3 Fw: TTAGGTGTGGCGCTCTCCAG. Rev: GCCAAGGACACGTCAGGATG; Hcn1 Fw: TGCCACAGCTTTGATCCAGT. Rev: GCGCATGTCAGCTGGTAACT; Kcnab1 Fw: TCCCATGGAAGAAATCGTTCG. Rev: CTTCCATGATCTCCATCGCG; Kcnd2 Fw: GGGAAGCCATAGAGGCAGTG. Rev: GTGGAGTTCTCTCCACACATCTG; Kcng3 Fw: CAACCGCAGTCTGGATGACC. Rev: CGGTGAACCACCCTATGCAG; Kcnh1: Fw: TTAGTGCCTTTATGGGTGATCC. Rev: GATGCCCTGACTCTCTCTCC; Kcnh5: Fw: CTTTCCAGACCCAGGATGCC. Rev: TTGATTCACTGGAGCGCCTG; Kcnj6 Fw: AACTGACGGAGAGGAATGGG. Rev: TGGGTTGGGTGAATTCTGGG; Kcnk2 Fw: ACGTGGCAGGTGGATCAGAC. Rev: AGTAGGCCAGCCCAACGAG; Kcnmb4 Fw: TTGGAAAGATGAGATCGGTTCC. Rev: AAGCAGTGCAGGAGAGCAATC; Kv2.1 Fw: CGTCATCGCCATCTCTCATG; Rev: CAGCCCACTCTCTCACTAGCAA; Gria1 Fw: CTGGTGGTGGTGGACTGTG. Rev: TGTCCATGAAGCCCAGGTTG; Gria2 Fw: ATTTGGAATGGTATGGTTGGAG. Rev: AGGCTCATGAATGGCTTCGAG; Gria3 Fw: TCATTCTCACGGAGGATTCCC. Rev: CACAGCAAAGCGGAAAGCAC; Gria4 Fw: CCAGAGCCTCCCTGAAGACC. Rev: TGCGCGCTCTCCTCTCTTTC; Grik1 Fw: AGTTGGTCGCATGCTCTTCG. Rev: CGAACCTTGATGCCCAATAC; Grik2 Fw: CTGGTGGAGAGTGCTTTGGG. Rev: CCACACCCTTGCAACCCTG; Grik3 Fw: TCGCCAGATTCAGCCCTTAC. Rev: GCCCATTCCAAACCAGAAGC; Grik5 Fw: GCCTGCGGTTGGTAGAGGAC. Rev: TCTGCCTTCCGGTTGATGAG; Grin1 Fw: TCAGAGCACACTGTGGCTGC. Rev: ATCGGCCAAAGGGACTGAAG; Grin2B Fw: GGAAGTGGGAGAGGGTGGG. Rev: CAGGACACATTCGAGGCCAC. For each reaction primers were used at 0.2μM using 10nl of total cDNA. Each sample was corrected to the housekeeping gene GAPDH.

### Dissection and western blot of mouse perinatal cortices

Postnatal pups (P7-8) were sacrificed by decapitation and fresh tissues were dissected immediately after. Whole cortex samples were collected and lysed in RIPA buffer (10 mM Tris-Cl (pH 8.0), 1 mM EDTA, 1% Triton X-100, 0.1% sodium Deoxycholate, 0.1% SDS, 140 mM NaCl, 1 mM PMSF) implemented with Complete protease inhibitors (Roche). Protein quantity was then measured with the Pierce BCA protein assay KIT (ThermoScientific). Equal amounts of proteins from tissue lysates were resolved by reducing SDS-PAGE, transferred to PolyVinylidene DiFluoride (PVDF) membrane and incubated with the indicated antibodies (see supplementary Table 2). Immunoblots were developed using ECL Prime western blotting detection reagent (GE Healthcare). Signals were normalized on actin signal intensity.

### Chromatin Immuno-Precipitation (ChIP)

The ChIP protocol was modified from Kuo and Allis 1999 (Methods), using protein sepharose-A resin (SIGMA), prepared by washing in PBS and equilibration buffer (HEPES pH7.4 20mM, EDTA 1mM, NaCl 150mM, Triton 0.8%, SDS 0.1% PMSF 10mg/ml). For each experiment (n=3), we dissected cortices of 8 mice at P0 in ice cold HBSS and washed in DMEM. Samples were crosslinked in 0.9 formaldehyde for 10’ and blocked adding 1.1ml of glycine 1.25M for 5’. We washed samples in wash buffer (HEPES pH 7.4 20 mM, NaCl 150mM, Glycine 0.125M, PMSF 10mg/ml) and isolated nuclei in Lysis buffer (HEPES pH 7.4 20mM, NaCl 150mM, Glycine 0.125M, SDS 1% PMSF 10mg/ml). Nuclei were collected (2200 rcf for 5’ spin) and sonicated in 1.2 ml Sonication Buffer (HEPES pH 7.4 20mM, NaCl 150mM, Glycine 0.125M, SDS 0.4% PMSF 10mg/ml) 6 times for 5 seconds at 10μm amplitude. DNA fragments were separated from cellular debris by spinning at 14000rcf for 10’ and recuperating the supernatant, fragmentation quality was evaluated by gel electrophoresis post-hoc. DNA fragments where split in 3x300 µl samples and precleared with resin Slurry solution (resin/equilibration buffer, 50/50 volume), adding 50ul of Slurry to each and incubating for 1h at 4°C. Supernatant was kept after spinning at 800 rcf for 2’ at +4°C and were incubated overnight at +4°C with 3µg of dedicated antibodies: Nr2f1 antibody (Thermo Fisher PA5-21611) and GFP antibody (Abcam ab13970, for aspecific binding). A fourth samples called Mock was obtained using a blank volume of Sonication buffer with no antibody. We added 50µl of Slurry to each sample and incubated 3h at +4°C and washed samples 4 times in ice cold Washing buffer A (HEPES pH 7.4 20 mM, NaCl 150mM, Triton 100 1%, SDS 0.1% PMSF 10mg/ml), 4 times in ice cold Washing buffer B (Tris pH 8.0 20mM, EDTA 1 mM, LiCl 250mM, NP-40 0.5%, Na-Deoxycholate 0.5%, PMSF 10mg/ml) and 2 times with TE buffer (Tris HCl pH 8.0 20mM, EDTA 1mM, PMSF 10mg/ml). We eluted DNA fragments denaturating antibodies adding 300μl of Elution buffer (NaHCO3 50mM, 1% SDS) and incubating in rotation 1h at RT. Input samples were generated by mixing 300μl of Elution buffer to 10μl of the supernatant from the first washing of a sample without antibody. We separated the fragment from the resin and antibodies by spinning twice at 8000 rcf for 2’ keeping the supernatant. Next, we reversed crosslinks adding 12μl of NaCl 5M and incubating overnight at +65°C. RNA was removed by adding 1μl of RNAse-A 10mg/ml and incubated for 30’ at +37°C; the reaction was blocked adding 6μl of EDTA 0.5M. Remaining proteins were digested adding 12μl of Tris HCl pH 8.0 and 1μl of proteinase K 10mg/ml for 1-2h at +45°C. Finally, samples were purified by phenol-chloroform extraction and precipitated into 20µl of H_2_Omq.

### qPCR of ChIP samples

qPCRs on ChIP was performed using KAPA Sybr Fast (Kapa Biosystems) according to manufacturer’s instructions. Samples were run on a LigthCycler II 480 (Roche) and the following primers were used: Nr2f1 binding site at Hcn1 locus Fw: TTTCTGCTTTCCCGTGAGAGC. Rev: AGCCTAGCCTAGGAAGCCAG; Nr2f1 binding site at Kcnab1 locus Fw: TGTGGTATCAGAGATGAAAGGC. Rev: AAGGGACCTCCCAGCTCTTC; Nr2f1 binding site at Kcnk2 locus Fw: CCCAAAGCTGGACTTGTGTAC. Rev: TTCCCCTGATCAAAATTCCATTC; Negative control (Egr1 binding site at Cacna1g locus) Fw: AGCACCAGCTCAGATCACTCC. Rev: CAGAAGAAACTGTGTCAAGAGC. For each reaction primers were used at 0.2µM using 0.5µl each sample. For proper quantification, a calibration curve was included in the reactions using 6 sequential 1:10 solutions of samples obtained from amplicon extraction of PCR performed on genomic DNA with each set of primers used. For every ChIP experiment the reaction was carried out on 3 replicates of specific antibody, unspecific antibody, no antibody, mock and input samples. The calculated concentration of every samples was normalized to the input, set to 1 and we then calculated the fold change between the samples with specific antibody against sample with unspecific antibody, which was normalized to 1. Finally, the fold change of specific/unspecific antibody samples was corrected by dividing for the fold change of specific/unspecific antibody samples obtained from the negative control primers.

### Statistical tests

Principal component analysis (PCA) was performed on R software using Factominer and the following parameters: V*rest*, R*in*, Rheobase, V*threshold*, APamplitude, APhalf-width, latency, Sag and ISI. In PCA plots, ellipses represent the 95% confidence and the square the center of mass for each cluster. Hierarchical clustering was performed with Euclidean distance, using both average burst frequency and correlation per condition. In Figure 2, we used 2 way ANOVA to determine relevant parameters driving differences for burst frequency, coordination and frequency variability, post-hoc testing was performed using Welch Two-samples test. In Figure 6A, we performed Welch’s unequal variances t-test analysis correcting the P-value for multiple testing using the Benjamini–Hochberg procedure, false discovery rate = 0.25. Data analysis scripts were custom-prepared in R (Computing, 2018) and MATLAB. Additional statistical analyses were also performed using Graphpad Prism7 and data are presented as mean ± SEM. *n* values represent biological replicates from ≥ 3 different brains and ≥ 3 different litters as stated in each figure legends or supplementary tables. No statistical tests were used to predetermine sample size but samples sizes were similar as reported in previous publications (Del Pino *et al*., 2017; Del Pino *et al*., 2013) or as generally employed in the field. No randomization was used. Whenever possible both genotypes were processed in parallel on the same day. *Ex vivo* electrophysiology experiments and analysis of all data were performed before genotyping. Differences were considered significant when P < 0.05. Data sets were tested for normality (Kolmogorov-Smirnov) or homoscedasticity (Leven test) before performing parametric (Student t-test, one-way or 2-way ANOVA followed by Bonferroni’s post-hoc test) or non-parametric tests (Welch’s unequal variances t-test or Mann-Whitney test) used to determine P-values.

## RESULTS

### Impaired intrinsic network and bursting activity in cultured *Nr2f1*-deficient cortical neurons

To understand whether spontaneous cortical network activity could be under the control of a transcriptional regulator known to act on somatosensory area patterning, we employed a conditional mouse in which *Nr2f1* is exclusively deleted from the cortex (referred to here as *Nr2f1cKO*). In this mouse line, thalamic differentiation is not affected and thalamic inputs reach cortical subplate cells around birth (Armentano *et al*., 2007). As a first step to measure intrinsically-generated cortical activity in the absence of subcortical inputs, we dissociated somatosensory cortical neurons at E18.5 and cultured them onto coated multielectrode arrays (MEA). This allowed us to follow activity in an *in vitro* embryo-derived model at different times of cortical network maturation and assess the possible effect of cell-autonomous developmental programs on functional network organization (**Figure1 and S1**).

Our recordings on primary cultured neuronal networks started at 9DIV, since 6DIV neuronal cultures failed to show any activity in either *Ctrl* or *Nr2f1cKO* conditions. A comparison of the average firing frequency of the earliest events taking place at 9DIV demonstrated a significant lower *Nr2f1cKO* network activity when compared to *Ctrl* (**Figure 1A, B**). While there was no difference in the average number of bursts (**Figure 1C**), the duration of these bursts was significantly increased in mutant neurons (cf. *Ctrl*, **Figure 1D**). Furthermore, firing activity was more clustered into bursts in mutant neurons (**Figure 1E**). Notably, the firing synchronization index was higher in *Nr2f1cKO* than in *Ctrl* (**Figure 1F**). Hence, loss of *Nr2f1* leads to an overall reduced firing frequency, but also to an increased tendency of firing in bursts rather than isolated spikes in 9DIV-derived cortical neurons. Together with the presence of longer burst duration and increased network synchronization, these data point to an involvement of *Nr2f1* regulating neuronal network excitability.

**Figure 1.**
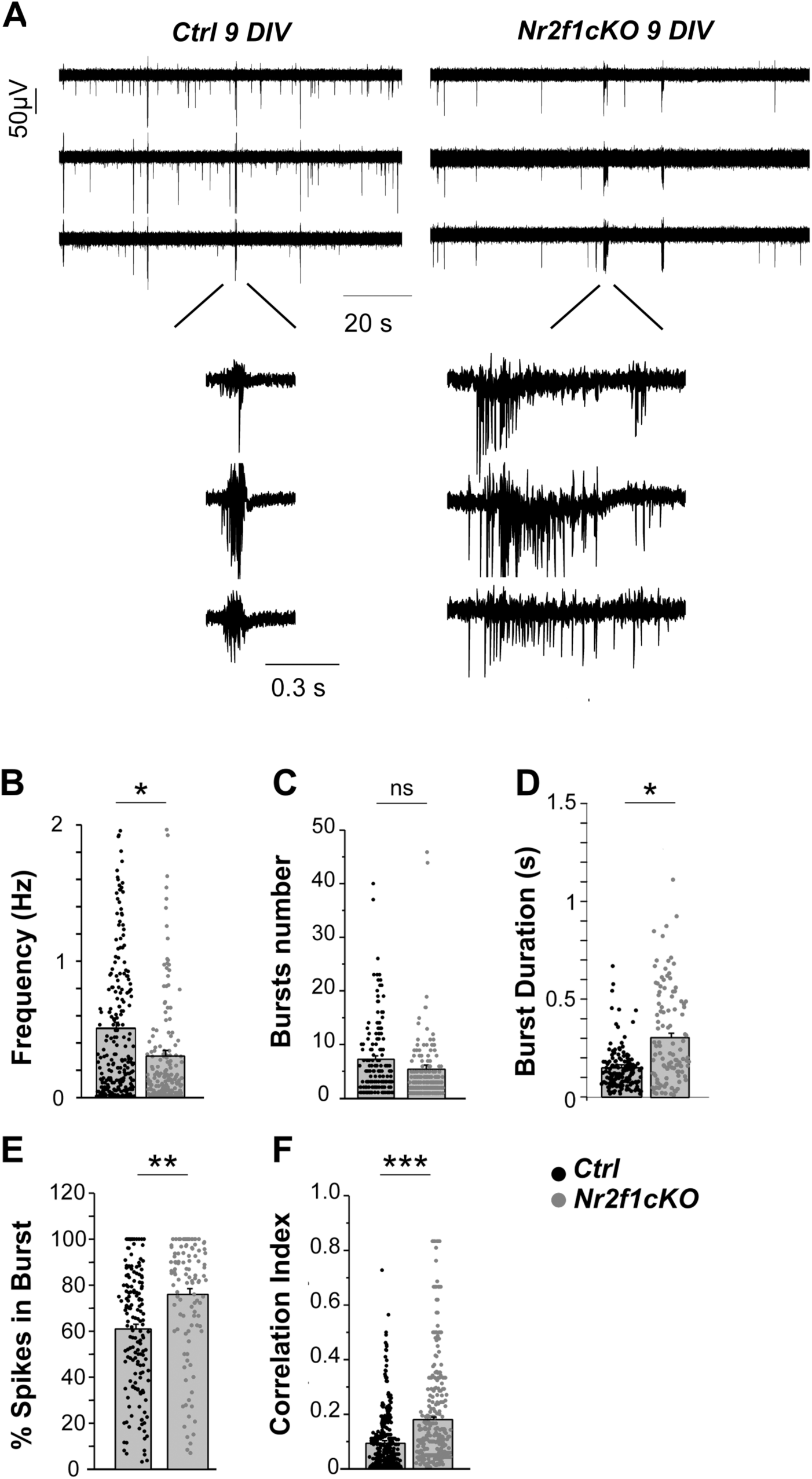
Impaired intrinsic network and bursting activity recorded in DIV 9 cultured Nr2f1-deficient cortical neurons. **A)** Spontaneous firing of *Ctrl* (left) and *Nr2f1cKO* (right) cortical network recorded by 3 representative MEA electrodes at 9DIV. In insets, representative bursts shown at expanded scale. **B-F**) Firing parameters in *Ctrl* (from N_MEA_=9) and *Nr2f1cKO* (from N_MEA_=7) are compared: frequency (*Ctrl*, 0.508 ±0.04, N_channels_= 271), *Nr2f1cKO*, 0.304 ± 0.042, N_channels_=208), bursts number (*Ctrl*, 7.211 ±0.67, N_channels_ = 170; *Nr2f1cKO*, 5.507 ± 2.726, N_channels_ =203), burst duration (*Ctrl*, 0.149 ±0.02, N_channels_ = 169; *Nr2f1cKO*, 0.306 ± 0.023, N_channels_ =112), % spikes in burst (*Ctrl*, 60.959 ±2.064, N_channels_ = 167; *Nr2f1cKO*, 76.029 ± 2.393, N_channels_ =114), cross correlation index (*Ctrl*, 0.1246 ±0.0007, N_channels_ = 256; *Nr2f1cKO*, 0.1453 ± 0.0009, N_channels_ =258), (* P<0.05; **P<0.01, unpaired T-test). Data are represented as mean ± SEM. **See also Figure S1.**

To further explore the maturation of this cortical network activity, we performed similar experiments in neuronal cultures at 15DIV and 20DIV (**Figure S1**). At 15DIV, we observed that the firing frequency of *Ctrl* cortical networks reached a maximum value, which then decreased at 20DIV (*data not shown*). We therefore compared firing parameters measured at 15DIV with those measured at 9DIV to assess whether network maturation was affected *in vitro*.

Interestingly, we observed that from 9DIV to 15DIV, the developmental increase in firing activity and in the number of bursts of somatosensory neurons was reduced in *Nr2f1cKO* neuronal networks (**Figure S1A-C**), together with the burst duration, which was significantly decreased in mutant neurons at 15DIV compared to 9DIV (**Figure S1D**). Finally, the percentage of spikes within bursts was also diminished in *Nr2f1cKO* (**Figure S1E**), in accordance with reduced cross-correlation probability index (**Figure S1F**). Since network maturation correlates with the expression of synchronized events, cross correlation probability between the firing at individual recording sites is representative of network maturation *in vitro* (Gavello *et al*., 2012; Gavello *et al*., 2018). Taken together, these results indicate that spontaneous network activity generated within the neocortex in the absence of extracortical inputs is affected upon loss of cortical *Nr2f1*. Thus, *Nr2f1* might be directly or indirectly involved in modulating intrinsic spontaneous network activity in the developing somatosensory cortex.

### Alterations in Ca2^+^ burst and synchronous activity in early postnatal *Nr2f1*-deficient cortices

As a further step to understand how spontaneous cortical network activity develops in the immature somatosensory cortex of *Ctrl* and *Nr2f1cKO* mouse pups, we set up a Ca^2+^ imaging protocol using the calcium indicator Oregon Green BAPTA^TM^ in *ex vivo* acute brain slices of P1 and P4 mouse pups. These two stages represent critical timepoints of early cortical somatosensory circuit establishment bridging the activity from the spontaneous thalamic waves (Anton-Bolanos *et al*., 2019) to the experience-driven thalamic inputs (Corlew, Bosma and Moody, 2004). Cells that displayed a bursting behavior typical of excitatory neurons were recorded over 15 minutes under standard perfusion (**Figure 2**). The time course of Ca^2+^ activity from individual cells was quantified using a moving average approach filtered for bursting cells, thus excluding inactive neurons (**Figure 2A**). Analysis of the Ca^2+^ burst frequency resulted in an overall increase from P1 to P4 in *Ctrl* (**Figure 2B**), in accordance with previous studies (Corlew, Bosma and Moody, 2004), but also in mutant slices. To evaluate the level of cortical network maturation, we measured the correlation index of Ca^2+^ bursts for each experiment, by quantifying the ratio of neurons participating to the same Ca^2+^ burst within a 3s window, and normalized across experiments on the total amount of bursts and cells. As expected, we observed increased correlation with network maturation progression (**Figure 2C**).

Next, we determined whether loss of Nr2f1 function would affect Ca^2+^ burst frequency and synchronization in the developing P1 and P4 cortices. Interestingly, we observed that Ca^2+^ burst frequency was higher in *Nr2f1cKO* brain slices than in the corresponding *Ctrl* cohorts at P1, but not at P4 (**Figure 2B**). Moreover, by comparing the Ca^2+^ burst synchronization events between *Ctrl* and *Nr2f1cKO* brains, we observed no differences at P1, but instead a decreased synchronization rate at P4 in mutant brains (**Figure 2C**). Since synchronization increases with time, these results suggest an impaired maturation of network complexity in mutant brains. Thus, although frequency of bursts and their correlation tend to increase in both conditions, the burst correlation increases more in *Ctrl* than in *Nr2f1cKO* brain slices, suggestive of defective maturation. Taken together, our data indicate that spontaneous network activity and synchronization patterns are altered upon loss of Nr2f1 function during prenatal stages.

**Figure 2.**
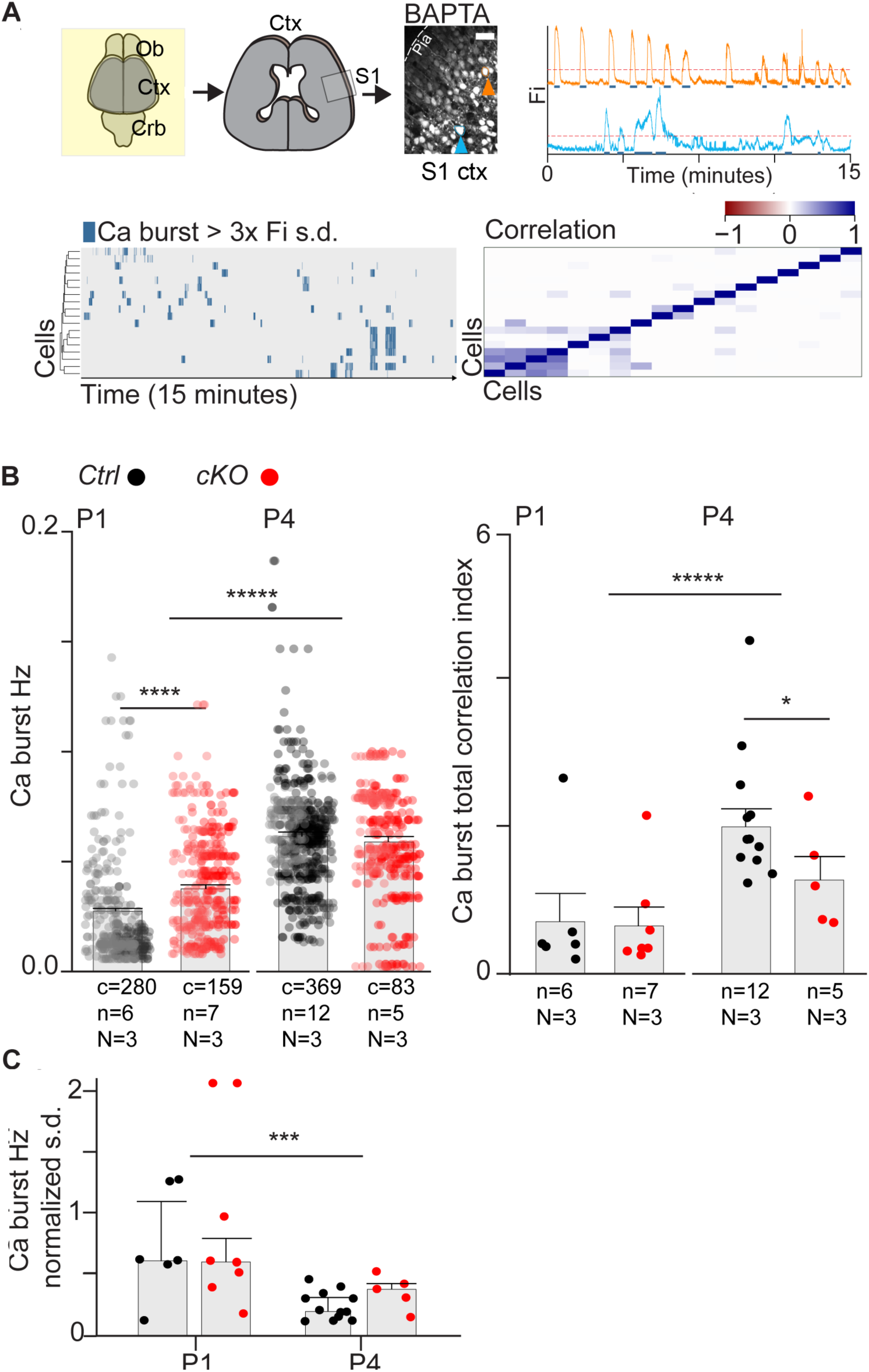
Altered Ca^2+^-dependent intrinsic activity upon Nr2f1 loss. **A**) Schematics of the acute horizontal slice preparation from somatosensory cortex and representative image of the Ca^2+^ BAPTA fluorescent signals. On the right, time series of intracellular signal in manually defined cells, corrected by moving average (Fi). Cells originating from the selected traces are indicated with colored arrows (orange for the upper and blue for the lower trace). Bursting events are defined by distinct thresholds. Below, time series matrices of above threshold Ca^2+^. Single traces have been sorted by Euclidean distance (left). Pearson correlation matrices of single neuron signal traces within single experiment (right). **B**) Distribution of single cell Ca^2+^ burst frequency (left) and burst intra-neuron correlation (right) in P1 and P4 brains. For Ca^2+^ bursts, single points indicate cells, for Ca^2+^ correlation, single points are single time-lapse images, each animal was recorded over more slices. We first used 2 way ANOVA to test frequency and correlation dependency on Genotype and Age, followed by post hoc analysis using Welch Two-samples test (P1 Ctrl n=6, P1 *Nr2f1cKO* n=7, P4 *Ctrl* n=12, P4 *Nr2f1cKO* n=5, Burst Frequency ANOVA: Genotype: FDR= 19.41, P=1.18e-05; Age: FDR=325.48, P=2e-16. Correlation ANOVA: Genotype: FDR=0.780, P=0.3849; Age: FDR=8.002, P= 0.0087. Whelch Two-samples post hoc test. P=0.8429. ****P=1.113e-8, *P= 0.04722, ***** P = 2.2e-16). **C**) Standard deviation of Ca^2+^ burst frequency normalized to the mean indicates a progressive decrease of frequency variability with maturation, while no substantial effect between genotypes is registered (2 way ANOVA: : Genotype: FDR= 0.231, P=0.063446; Age: FDR= 12.798, P=0.00134. Whelch Two-samples post hoc test P=0.00679 ***). Statistical scripts were custom-prepared in R (Computing, 2018). Abbreviations: Ob, Olfactory bulb; Ctx, Cortex; Crb, Cerebellum; S1, somatosensory cortex. N, Number of animals; n, number of slices; c, number of cells. Scale bar: 10µm.

### Abnormal intrinsic electrophysiological properties in early postnatal *Nr2f1*-deficient cortical layer V pyramidal neurons

As intrinsic electrophysiological properties are major determinants of spontaneous neural activity, we next examined the state of the intrinsic excitability of layer V glutamatergic pyramidal neurons (LVPNs) in *Nr2f1cKO* mouse cortices, previously shown to express an altered molecular signature (Tomassy *et al*., 2010; Harb *et al*., 2016). We measured their intrinsic excitability profile in acute brain slices from *Ctrl* and *Nr2f1cKO* mouse pups during postnatal day (P)5–P8, when corticogenesis and neural migration already took place. We employed whole-cell recordings in current-clamp mode, and performed the measurements in the absence of synaptic blockers since, at this stage, the number of synaptic inputs resulted very low, in agreement with the low number of synaptic connections reported in other studies (Favuzzi *et al*., 2019). In order to compare electrophysiological properties of *Ctrl* and *Nr2f1cKO* neurons, we aimed to target similar populations by selecting neurons that were identified as LVPNs through *post-hoc* biocytin-filling, immunohistochemistry and morphological reconstruction. Although LVPNs differentiate into two major types in terms of morphology and intrinsic physiology in adult brains, these two LVPNs types are not well differentiated at the electrophysiological level during early postnatal stages (Kasper *et al*., 1994; Christophe *et al*., 2005). Among all recorded cells, we analyzed 19 *Ctrl* and 28 *Nr2f1cKO* neurons (from 8 and 7 different brains, respectively) located in layer V and displaying a slowly adapting regular spiking firing pattern (**Figure 3A**). To first explore physiological differences between *Ctrl* and *Nr2f1cKO* LVPNs, we performed multivariate statistical analysis from intrinsic electrophysiological parameters. This revealed that *Nr2f1cKO* LVPNs clearly segregate from the *Ctrl* population (**Figure 3B**). Notably, specific changes in passive and active electrophysiological properties were observed in *Nr2f1cKO* neurons. Mutant LVPNs displayed a significantly more depolarized resting membrane potential and therefore a decreased rheobase when compared to *Ctrl* LVPNs. Interestingly, *Nr2f1cKO* LVPNs also exhibited a significantly reduced voltage sag in response to hyperpolarizing current steps, suggestive of reduced I_h_ currents compared to *Ctrl* LVPNs (**Figure 3C**). No significant changes in input resistance, action potential (AP) threshold, AP amplitude, AP half-width, inter-spike interval, membrane time constant or input/output function were observed in mutant LVPNs (**Figure S2, Table S1**). Taken together, this functional characterization suggests that Nr2f1 deficiency in the developing cortex leads to specific alterations in intrinsic excitability of immature LVPNs in the somatosensory region.

**Figure 3.**
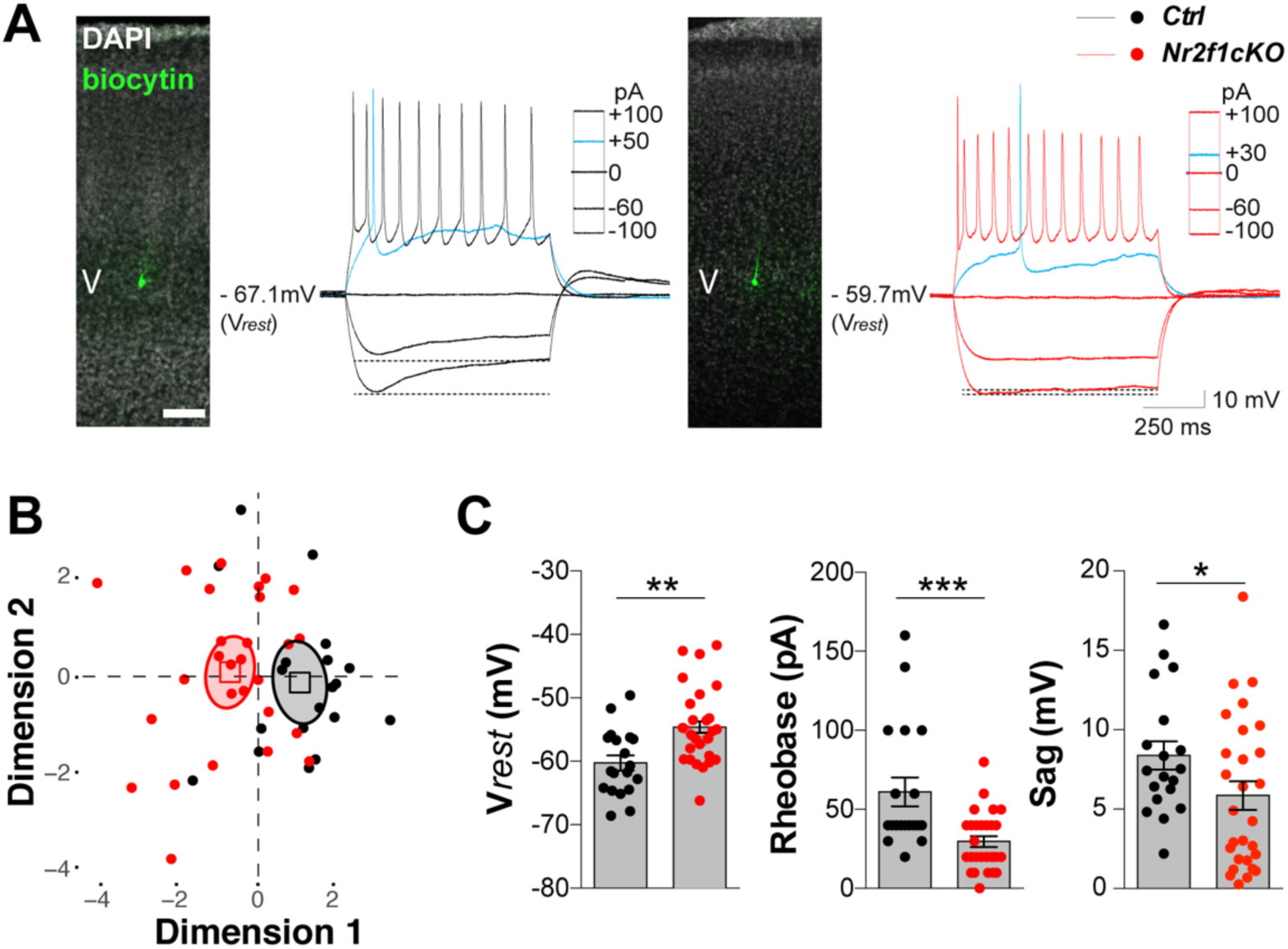
Abnormal intrinsic excitability of Nr2f1-deficient layer V pyramidal neurons. **A**) Layer V pyramidal neurons (LVPNs) identified post-hoc through fluorescent labelling for biocytin (green) and DAPI (gray) in *Ctrl* (in black) and *Nr2f1cKO* (in red) somatosensory cortex display slow-adapting regular spiking firing pattern. **B**) Individual factor maps (PCA) performed on intrinsic electrophysiological properties of LVPNs segregate two clusters as distinct populations in *Ctrl* (black) and *Nr2f1cKO* (red). **C**) The mean resting membrane potential (V*rest*) is significantly more depolarized (\*\**P* = *0.0013*), while the Sag (\**P* = 0.0418) and rheobase (\*\*\**P* = *0.0010*) are significantly decreased in *Nr2f1cKO* LVPNs compared to *Ctrl* LVPNs. (N = 19 *Ctrl*, N = 28 *Nr2f1cKO*, from 8 and 7 different brains respectively; two-tailed unpaired T-test or U Mann-Whitney test). V, cortical layer V. Data are represented as mean ± SEM. Scale bar: 100 µm. **See also Figure S2 and Table S1.**

### Reduced dendritic complexity of LVPNs in *Nr2f1* mutant cortices

To assess whether abnormal intrinsic excitability of mutant LVPNs was accompanied by changes in dendritic morphology, we reconstructed and analyzed the morphology of previously recorded LVPN filled with biocytin (**Figure 4A,B**). We first evaluated dendritic arbor geometry and complexity by performing Sholl analysis (**Figure 4C, S3 and Table S2**). Comparison of Sholl profiles revealed that *Nr2f1cKO* LVPNs have a significantly reduced proximal dendritic branching pattern (20-100µm from the soma) when compared to *Ctrl* LVPNs, whereas minor or no differences were observed in distal regions (**Figure 4, S3A and Table S2**). Next, to evaluate more specific features of dendrite geometry, we assessed the basal dendrite complexity index (DCI), arbor length, number of primary dendrites, branching points and tips, and oblique formations of basal and apical dendrites (**Figure 4D-H and S3B-D**). Our morphological data point to a less elaborate basal dendrite complexity in mutant LVPNs (**Figure 4 and S3D**), whereas no significant differences were observed in apical and oblique dendrite geometry between *Ctrl* and *Nr2f1cKO* neurons. In particular, the basal DCI was significantly reduced in *Nr2f1cKO* compared to *Ctrl* LVPNs (**Figure 4D–E**), as a result of a reduced number of primary dendrites, branching points and terminal tips (**Figure 4F–H**). Overall, our data indicate that loss of Nr2f1 function affects the complexity of basal dendrite geometry in LVPNs at early postnatal stages.

### Reduced conformation and plasticity properties of the axon initial segment in mutant *Nr2f1* LVPNs

Intrinsic excitability and dendritic changes are often accompanied by variations in structural properties of the axon initial segment (AIS), a specialized structure playing a pivotal role in integration of synaptic inputs and action potential initiation (Yamada and Kuba, 2016; Kole and Stuart, 2012). It has been reported that distinct AIS features, such as length and/or location relative to the soma, are crucial for the homeostatic control of neuronal excitability in layer V pyramidal neurons (Hamada *et al*., 2016). The observed reduction of basal dendrite complexity, sag response and the presence of a more depolarized *Vrest* together with the lack of effect on the input-output relationship (**Figure 3 and S2**), led us to hypothesize that structural homeostatic scaling of the AIS could occur in *Nr2f1cKO* LVPNs. This could act as a potential mechanism stabilizing action potential generation, as observed in previous studies (Hamada *et al*., 2016; Gulledge and Bravo, 2016). To test this hypothesis, we analyzed AIS structure of P7 LVPNs *Ctrl* and *Nr2f1cKO* somatosensory cortices employing the cytoskeletal scaffolding protein Ankyrin G – a widely used marker of the AIS – (Kordeli, Lambert and Bennett, 1995; Zhou *et al*., 1998), together with CTIP2, a layer V marker (Arlotta *et al*., 2005), and NeuN, a neuronal marker labelling the soma (**Figure 5**). Notably, we found a significantly reduced length and diameter of the Ankyrin-positive AIS in *Nr2f1cKO* compared to *Ctrl* LVPNs (**Figure 5A–C**), indicating that altered intrinsic excitability and structural dendritic changes of mutant LVPNs are accompanied with structural changes in AIS conformation (Grubb and Burrone, 2010; Wefelmeyer, Puhl and Burrone, 2016; Kuba, Oichi and Ohmori, 2010; Kuba, 2010). Decreased length and diameter were maintained in P21 LVPNs (**Figure 5D-F**), suggesting that altered AIS structure observed at P7 is not the result of a delayed neuronal maturation.

**Figure 4.**
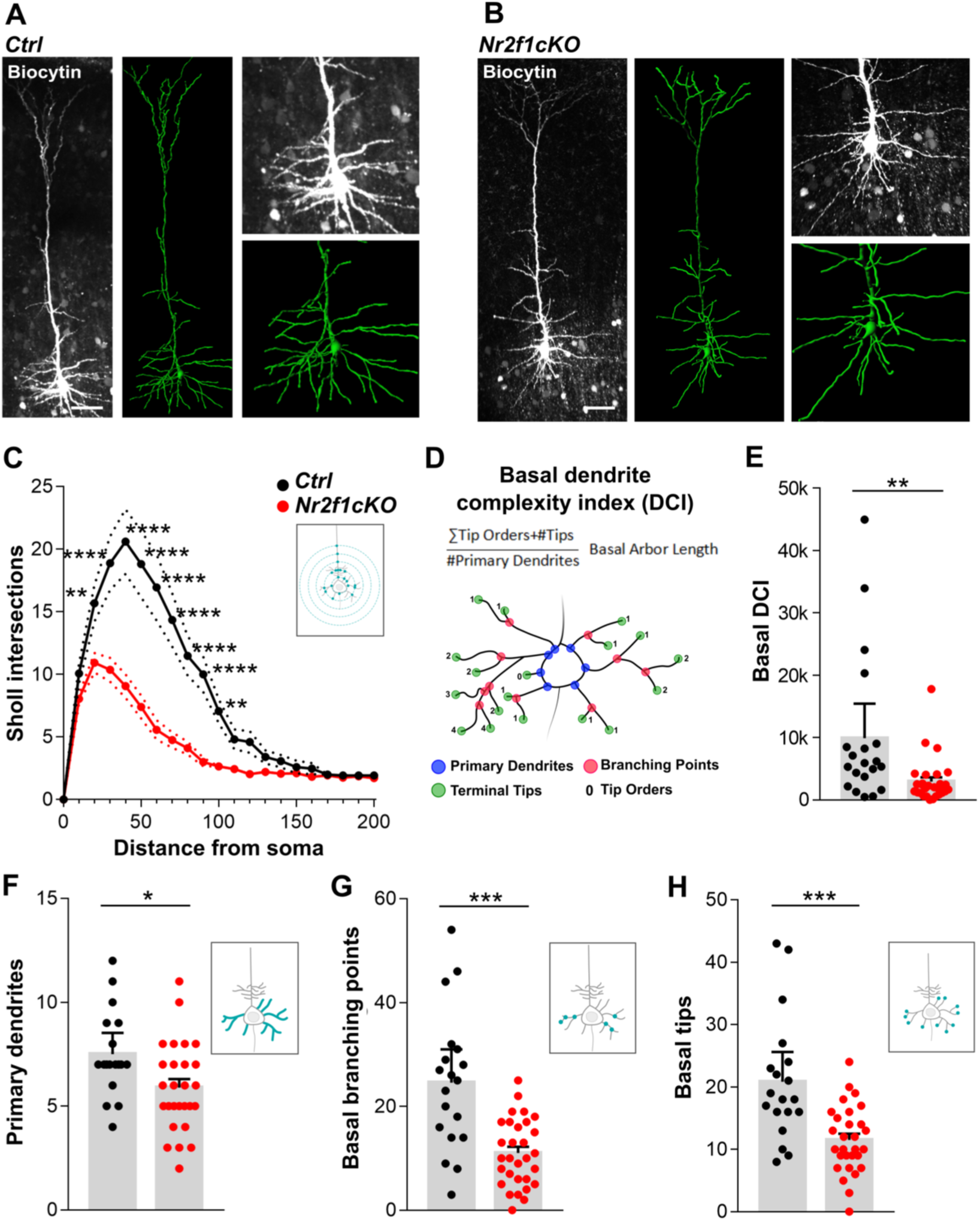
Reduced basal dendritic complexity of layer V pyramidal neurons (LVPNs) in Nr2f1 mutant cortices. Representative images of *Ctrl* **(A)** and *Nr2f1cKO* **(B)** LVPNs. In grayscale, maximum intensity projection of a multi-stack confocal image; in green, correspondent 3D reconstructions. On the right, magnifications of the basal structures. Scale bar: 50μm. **C)** Sholl analysis points to a reduction of the arborization complexity of *Nr2f1cKO* LVPNs (\*\**P* = *0.002*; \*\*\*\**P* < *0.0001*). **D**) Basal Dendrite Complexity Index (DCI) formula and schematic representation of primary dendrites, branching points, terminal tips and tip orders. **E-H**) Average basal DCI (*Ctrl*, 9902 ± 2650; *Nr2f1cKO*, 2956 ± 658; *P* = *0.0019*), number of primary dendrites (*Ctrl*, 7.5 ± 0.49; *Nr2f1cKO*, 5.89 ± 0.41; P = 0.0159), basal dendrites branching point (*Ctrl*, 24.58 ± 3.05; *Nr2f1cKO*, 11.03 ± 1.17; *P* = *0.0004*) and terminal tips (*Ctrl*, 20.84 ± 2.265; *Nr2f1cKO*, 11.52 ± 0.99; *P* = *0.0009*) are statistically reduced in mutant cells compared to controls. (N = 19 *Ctrl*, N = 30 *Nr2f1cKO*. N defined as number of analyzed cells). P-values are calculated by 2way ANOVA test **(C)**, Mann-Whitney non parametric test **(E)** or Welch’s unequal variances t-test **(F-H).** Data are represented as mean ± SEM. **See also Figure S3 and Table S2.**

**Figure 5.**
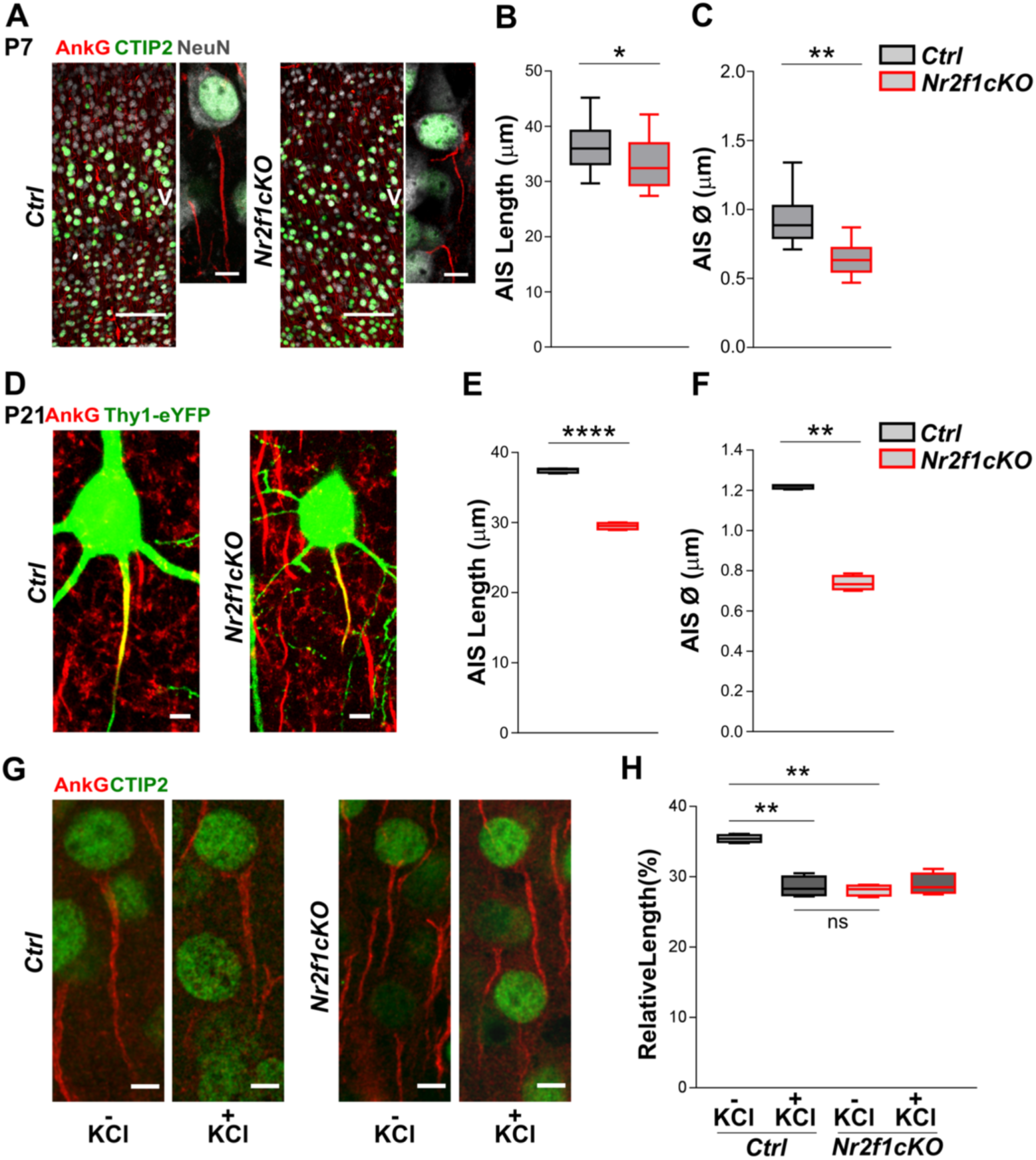
Abnormal structural features of the axon initial segment (AIS) in mutant Nr2f1 layer V pyramidal neurons. **A**) Left panel, coronal sections of P7 somatosensory *Ctrl* and corresponding *Nr2f1cKO* cortices labelled with CTIP2, for layer V identification, Ankyrin G for the AIS and NeuN as a somatic marker. Scale bar: 100 μm. Right panel confocal images of triple stained *Ctrl* and *Nr2f1cKO* LVPNs. Scale bar: 10 μm. **B-C**) Average AIS length (*Ctrl*, 35.23 ± 0.67; *Nr2f1cKO*, 31.14 ± 0.32; *P=0.0129*) and diameter (*Ctrl*, 0.94 ± 0.006; *Nr2f1cKO*, 0.65 ± 0. 004; *P=0.0043*) are statistically reduced in *Nr2fcKO* neurons compared to *Ctrl* at P7. (N=3; N defined as number of distinct analyzed animals). **D**) Confocal images of *Ctrl* (left) and *Nr2f1cKO* (right) LVPNs from P21 somatosensory cortices. In green *Thy1-eYFP* reporter gene, in red Ankyrin G. **E-F**) Average AIS length (*Ctrl*, 37.46 ± 0.15; *Nr2f1cKO*, 29.51 ± 0.23; *P<0.0001*) and diameter (*Ctrl*, 1.2 ± 0.005; *Nr2f1cKO*, 0.74 ± 0.02; *P<0.0001*) are statistically reduced in *Nr2f1cKO* cells compared to controls at P21. (N=3; N defined as number of distinct analysed animals). **G**) Confocal images of control and *Nr2f1cKO* LVPNs non-treated (-KCl) and treated (+KCl) with high KCl aCSF. In green CTIP2, in red Ankyrin G. **H**) Average relative length (average length values were normalized on control values) are significantly reduced in controls after KCl treatment (*Ctrl* – *KCl*, 35.41 ± 0.26; *Ctrl* + *KCl*, 28.57 ± 0.71; *P<0.0001*). No differences were observed in *Nr2f1cKO* cells between non-treated and treated populations. (*Nr2f1cKO* – *KCl*, 28.1 ± 0.3756; *Nr2f1cKO* + *KCl*; 28.9 ± 0.7714; *P=0.289*). (N=3; N defined as number of distinct analysed animals). P-values are quantified by the Welch’s unequal variance t-test (**B**), Unpaired t-test (**C-F**) or 2way ANOVA test (**H**).

Finally, to assess whether *Nr2f1*-deficient LVPNs would still respond to depolarizing input despite their reduced AIS length and diameter, we performed chronic depolarization treatment (+7mM of KCl for 6h) in acute brain slices of *Ctrl* and *Nr2f1cKO* P7 pups. In agreement with previous studies (Dumitrescu, Evans and Grubb, 2016), the depolarization treatment reduced the relative AIS length of *Ctrl* LVPNs. In contrast, no significant changes were observed in AIS size between depolarization and control treatment of *Nr2f1*-deficient LVPNs (**Figure 5G, H**). These data indicate that mutant LVPNs are not able anymore to undergo AIS plasticity upon increases in network activity.

### Altered expression of ion channels and glutamate receptors in postnatal cortices of *Nr2f1cKO* mutants

Since neuronal intrinsic excitability and AIS functionality of immature neurons are primarily determined by the expression, subcellular distribution and biophysical properties of ion channels, we next performed RT-qPCR for an extensive list of K^+^, Na^+^ and Ca^2+^ channels specifically expressed in deep layers, from *Ctrl* and *Nr2f1cKO* cortical lysates (**Figure 6A**). Out of 33 ion channels that we could successfully test, we found a significant reduction in the *mRNA* levels of only *Hcn1*, *Kcnab1*, *Kcnk2*, *Cacna2d1*, *Gria3* and *Grin1* in *Nr2f1cKO* when compared to *Ctrl* brains (**Figure 6A and S4**). Notably, we found significantly decreased protein levels of the hyperpolarization-activated cation channel HCN1 (Shah, 2014) in *Nr2f1cKO* when compared to *Ctrl* cortices (**Figure 6B, S4 and S5**). Moreover, double staining of HCN1 and CTIP2 on cortical sections at P7, when HCN1 expression becomes detectable in cell bodies (**Figure 6C**), confirmed decreased HCN1 protein signal intensity in CTIP2-expressing LVPNs, as well as a reduced number of double CTIP2/HCN1 positive neurons (**Figure 6D,E**), supporting reduction of HCN1 protein in layer V neurons of *Nr2f1cKO* mutant brains.

**Figure 6.**
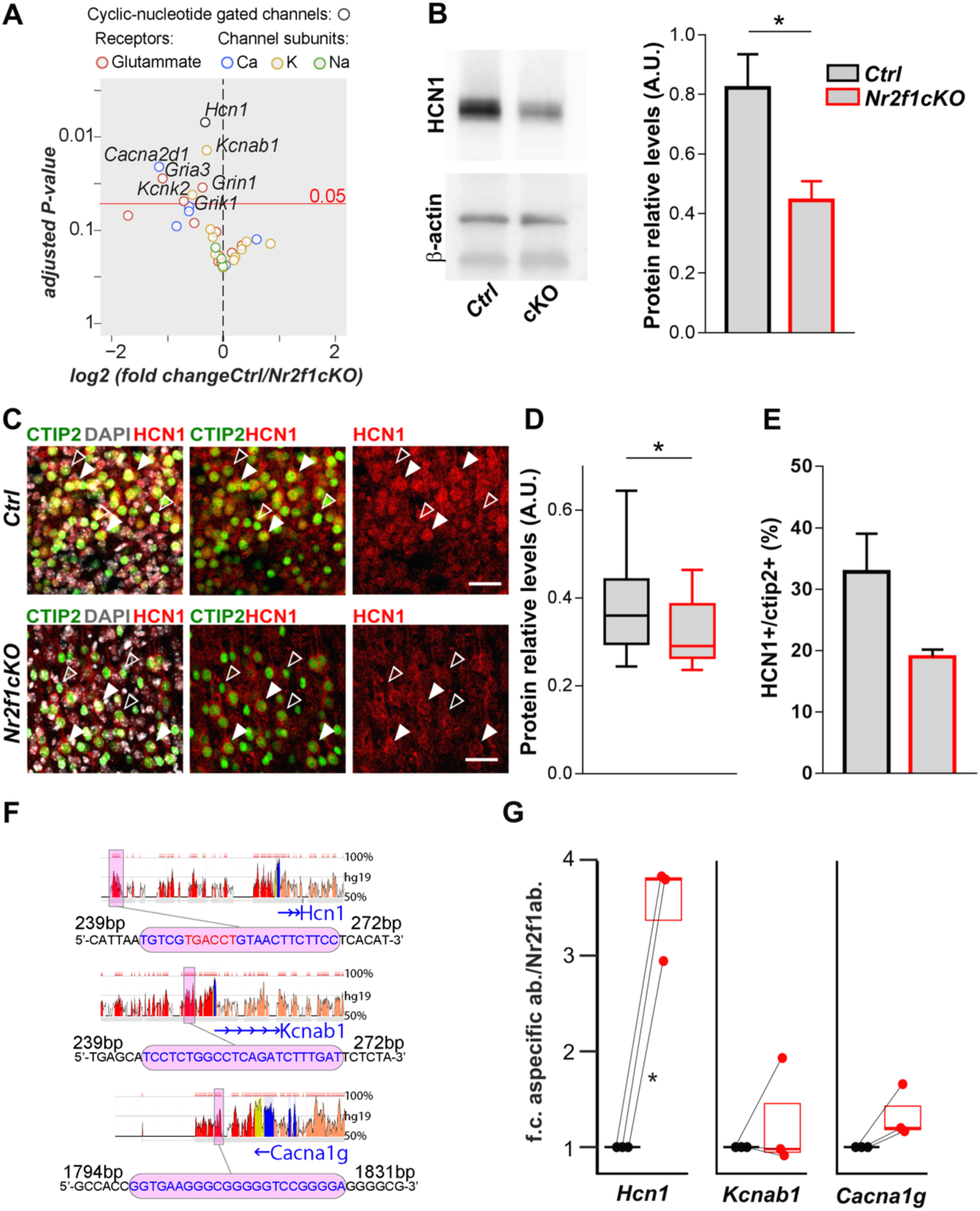
Regulation of ion channel-expression levels by Nr2f1. **A**) Data plot of qPCR analysis on P7 *Ctrl* and *Nr2f1cKO* P7 cortices (N≥3 brains). The red line represents the threshold for significance. Significantly downregulated genes are depicted in the upper-left panel. **B**) Western- blot analysis of HCN1 protein shows a significant reduction in *Nr2f1cKO* P7 cortices (*Ctrl*, 0.822 ± 0.11;*Nr2f1cKO*,0.44 ± 0.064; *P=0.05*; N=4). **C**) CTIP2-HCN1 double IF staining of P7 somatosensory cortices. **D**) Box and whiskers plot distribution of HCN1 cell signal intensity points to a reduction of HCN1 in *Nr2f1cKO* LVPNs (*Ctrl*, 0.89 ± 0.03; *Nr2f1cKO*, 0.71 ± 0.01; *P* = 0.015; N = 3). **E**) The number of HCN1+ cells within the LVPN CTIP2^+^ population tends to be reduced in Nr2f1cKO cells (*Ctrl*, 32.81 ± 6.24; *Nr2f1cKO*, 18.99 ± 1.19; *P = 0.15*; N = 3 brains). **F**) Snapshots from Evolutionary Conserved Regions (ECR) browser showing Nr2f1 binding location and sites (in pink; red nucleotides highlighting the canonical direct repeat of *Nr2fI* binding site: TGACCT); intergenic regions (red); UTR (yellow); exons (blue); introns (orange). **G**) Graph showing the ratio between unspecific and specific antibody (a.b.) fold changes (f.c.) on Nr2f1 consensus sites. N=3. P-values are calculated by Welch’s unequal variances t-test (A-E), or paired (G) t-test followed by Benjamini–Hochberg multiple test correction, false discovery rate = 0.25 (A). Data are represented as mean ± SEM. **See also Figure S4 and Figure S5**.

Finally, we investigated whether voltage gated ion channels whose transcript levels were found altered in mutant brains, were directly regulated by Nr2f1. To address this, we searched for evolutionary conserved binding sites for Nr2f1 on the group of differentially-expressed genes using the ECR browser tool (Loots and Ovcharenko, 2007; Ovcharenko *et al*., 2004). By using MatInspector (Quandt *et al*., 1995) and ChromAnalyzer (Montemayor *et al*., 2010), we identified putative binding sites for Nr2f1 at 25kb of the 5’UTR of *Hcn1*, 3kb of the 5’UTR of *Kcnab1* and proximal to the 5’UTR of *Kcnd2* loci (**Figure 6F**). These sites mainly match the Nr2f1 canonical binding sequence consisting of direct repeats with variable spacer configurations, as previously described (Montemayor *et al*., 2010). Notably, we observed that the binding sequence on the *Hcn1* locus matched the expected consensus site (**Figure 6F**, red nucleotides). To experimentally test direct regulation of the expression of these ion channels by Nr2f1, we performed chromatin immunoprecipitation (ChIP) assay on whole newborn *Ctrl* cortices using a well-established Nr2f1 antibody (Alfano *et al*., 2011; Parisot *et al*., 2017). Quantitative PCR (QPCR) analysis of immunoprecipitated material showed a reproducible 3- to 4 -fold significant enrichment of Nr2f1 binding *versus* GFP non-unspecific binding only on the *Hcn1* locus and not on the *Kcnab1* or *Cacna1g* loci (**Figure 6G**), despite their altered transcript levels quantified in mutant brains (**Figure S4**). These data on the molecular landscape of the somatosensory cortex indicate that while expression levels of several ion channels might be modulated by Nr2f1 during cortical development, only few of them seem to be directly controlled by Nr2f1. We identified *Hcn1* gene as one of these putative direct targets, whose protein product is known to strongly determine voltage sag and intrinsic excitability (Brager, Akhavan and Johnston, 2012; Fan *et al*., 2016), in agreement with our electrophysiological data (**Figure 3**).

## DISCUSSION

Our work reveals that the area patterning gene *Nr2f1* can influence the emergence of spontaneous activity and intrinsic excitability within the immature somatosensory cortex during early postnatal development. Our findings reinforce the role for *Nr2f1* in somatosensory area specification, as previously reported (Alfano *et al*., 2014; Armentano *et al*., 2007). More importantly, they provide, to our knowledge, first experimental proof for a transcriptional regulator, involved in cortical map formation, in determining directly or indirectly early patterns of spontaneous network activity intrinsic to the immature cortex. Finally, our data demonstrate a primary link between intrinsic genetic determination and spontaneous cortical activity through the regulation of voltage-gated ion channels in pyramidal glutamatergic neurons (summarized in **Figure 7**). We propose that, among others, the hyperpolarization-activated cation channel HCN1 – known to contribute to intrinsic neural excitability and spontaneous rhythmic activity in the brain (Shah, 2014)– might represent one key player during the establishment of neuronal synchronicity in the developing neocortex.

**Figure 7.**
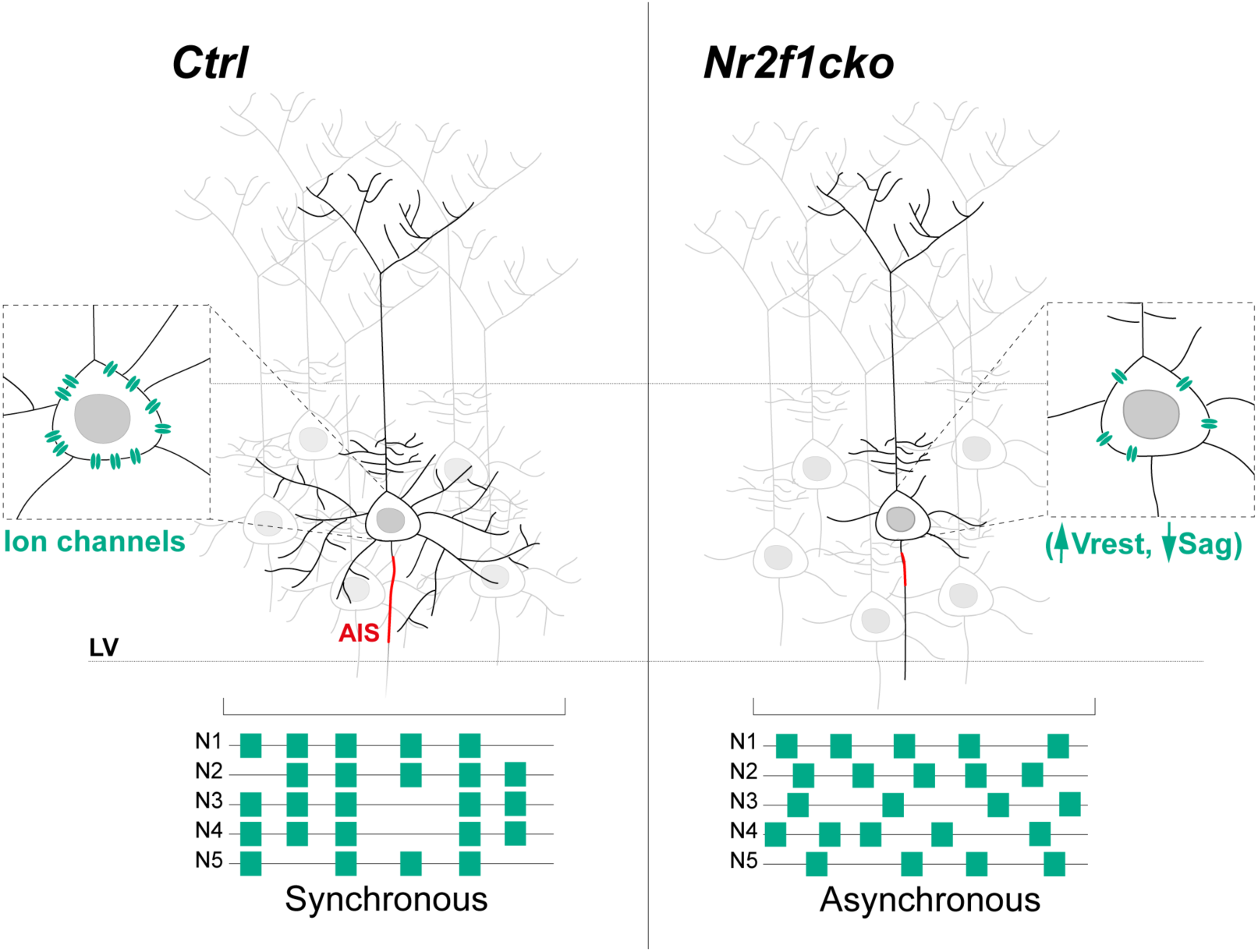
Summary schematics of changes in pyramidal neuron anatomy, bioelectric properties and network synchronization upon Nr2f1 loss. The summary schema highlights a pyramidal neuron with less complex basal dendritic arbor, shorter AIS (axon initial segment, in red) and reduced expression of ion channels (in green), leading to altered resting membrane potential (Vrest) and sag ratio (Sag) as well as asynchronous firing of neurons (below) within the cortical network of the immature somatosensory area, in the absence of *Nr2f1* (*Nr2f1cKO*) compared to a *Ctrl* neuron.

### Cortical and thalamic prenatal activity governing somatosensory maps

Developmental spontaneous activity has already been described to follow a regionalized pattern in the cortex (Corlew, Bosma and Moody, 2004; Uhlen *et al*., 2015) and to ultimately influence the assembly of long range circuits (Tritsch *et al*., 2007; Blankenship *et al*., 2011; Yamamoto and Lopez-Bendito, 2012; Schneggenburger and Rosenmund, 2015; Anton-Bolanos *et al*., 2019). However, how spontaneous activity is first triggered and then finely modulated during formation of the neocortical protomap is still unclear. In this study, genetic manipulation targeting exclusively cortical neurons has allowed us to investigate the specific implications of abnormal somatosensory area patterning for intrinsic network activity within the immature neocortex. We showed that an area patterning gene known to determine sensory identity of early differentiating neurons (Alfano *et al*., 2014) can contribute in establishing intrinsic network activity within the neocortex by regulating voltage-gated ion channels in pyramidal glutamatergic neurons. A recent study shows that prenatal activity from thalamic neurons can also influence the formation of a somatosensory map in the cortex before the emergence of sensory inputs (Anton-Bolanos *et al*., 2019). In the absence of thalamic calcium waves, the neocortex becomes hyperexcitable, the columnar and barrel organization are perturbed, and the somatosensory map lacks anatomical and functional structure. This could suggest that impaired formation of the somatosensory map observed in our *Nr2f1cKO* mutants might be caused by the absence of prenatal thalamic inputs reaching the mutant cortex. We exclude this possibility, since thalamic axons do reach the subplate in *Nr2f1* mutant prenatal cortices (Armentano *et al*., 2007). Hence, spontaneous activity intrinsic to the neocortex and the thalamus are both necessary for the functional assembly of somatotopic maps. Our data suggest that the neocortex needs to be electrically competent to receive thalamic waves at perinatal stages, and that early expression of arealization genes, such as Nr2f1, might be necessary to set up proper and synchronized activity patterns within the immature somatosensory cortex.

### Determinants of spontaneous network activity controlled by area patterning genes

Normal developmental trajectories in the cerebral cortex are characterized by a sequential process of spontaneous activity patterns in immature networks. In the mouse neocortex, large plateaus of synchronized network activity at birth are followed by early network oscillations and large depolarization potentials along the first postnatal week (Allene *et al*., 2008; Allene and Cossart, 2010). Multiple aspects that relate to intrinsically regulated maturation of pyramidal neurons might influence the developmental sequence of cortical network activity. Our data provide a novel characterization at the electrophysiological and molecular levels of intrinsic excitability features directly or indirectly regulated by the transcriptional regulator *Nr2f1* during the first postnatal week. Our conclusion that *Nr2f1* expression influences the molecular determinants relevant for early neuronal network activity is corroborated by previous transcriptome analyses, in which *Nr2f1* loss in post-mitotic cortical cells regulate the expression of several activity related genes, such as *Chrna7, Kcnq3, Kcnlp4, Robo1* and *Sv2b* (Alfano *et al*., 2014). However, the direct impact of these candidates on immature cortical neuron activity still remains to be investigated.

Other recent studies also highlighted the role of cell intrinsic and pre-synaptic factors in determining cortical network development with potential long-term effects (Boillot *et al*., 2016; Murase *et al*., 2016). In addition, a recent report has shown that the transcription factor Tbr1, known to regulate cortical layer identity, also alters the intrinsic excitability of neocortical neurons, by regulating HCN1 transcription in neonates and increasing sag response in layer VI (Fazel Darbandi *et al*., 2018). As Tbr1 is also a key factor for cortical development, it is presumably regulating spontaneous network activity intrinsic to the neocortex during early developmental stages, but this still remains to be addressed. Thus, regulation of HCN1 levels might be one of many other ion channels used as common mechanism by early transcriptional regulators to modulate spontaneous network activity within the developing cortex.

Within immature cortical networks, the earliest patterns of correlated spontaneous activity – e.g. synchronous plateau assemblies – are thought to arise from voltage-gated conductance and gap junction coupling (Pearson *et al*., 2004; Roerig and Feller, 2000; Blankenship *et al*., 2011; Cho and Choi, 2012). Our work features intrinsic excitability – more precisely, intrinsic bioelectrical properties of neural membranes – as an important component of neocortical development, likely contributing to the abnormal cortical patterning defects displayed in *Nr2f1* mutants. In this context, it is important to note that *Nr2f1* loss also affects the expression of many cell-adhesion related genes *(i. e., Cdh4, Cdh11, Cdh12, Fat3,* and *Ncam2*) (Alfano *et al*., 2014), which could in turn affect gap-junction functionality. Interestingly, intrinsic bioelectrical membrane properties prominently determine the occurrence of membrane potential spikes before gap-junction coupling takes place (Allene and Cossart, 2010). Hence, it is tempting to speculate that even earlier forms of activity within the embryonic brain could be responsible for the abnormal area specification observed in the absence of *Nr2f1* (Armentano et al., 2007; Alfano et al., 2014).

### Nr2f1-dependent *versus* homeostatic changes in intrinsic neural excitability

Our functional analysis indicates that an abnormally differentiating somatosensory cortex displays increased spontaneous network activity at P1 followed by decreased synchronization within immature neurons at P4. Principal factors for increased excitability of neural network might be the more depolarized resting membrane potential of pyramidal glutamatergic neurons, which might contribute to the higher frequency of Ca^2+^-events observed at P1. Alternatively, a decreased expression of voltage sag (I_h_) could lead to opposite changes in membrane excitability, which are strongly age-dependent (Bender and Baram, 2008; Magee, 1998; Williams and Stuart, 2000; Williams and Stuart, 2003; Berger, Larkum and Luscher, 2001; Poolos, Migliore and Johnston, 2002; Poolos, Bullis and Roth, 2006; Wang *et al*., 2003; Fan *et al*., 2005; van Welie *et al*., 2006; Brager and Johnston, 2007; Chen *et al*., 2001). Multiple membrane ion channels, whose expression is altered in pyramidal neurons upon loss of *Nr2f1* function, could also contribute to the increase in resting membrane potential in immature neurons, a property of pyramidal neurons that matures during the two first weeks after birth. In our *ex vivo* electrophysiological analysis, some membrane properties of *Nr2f1cKO* LVPNs, such as *Vrest*, resemble those of less mature neurons. However, other electrophysiological parameters, e.g. R*in*, do not change and therefore are not in agreement with a delayed maturation in intrinsic properties. In addition, our analysis on AIS homeostatic plasticity at P21 does not point to a maturation defect in individual neurons, but rather to a stabilized high state of abnormal intrinsic excitability leading to decreased dendritic complexity and AIS plasticity, as a possible homeostatic/compensatory mechanism.

Differences in ion channel transcripts support alterations in the endogenous activity of developing sensory areas, most probably affecting the overall frequency at early stages and leading to asynchronized network activity at later stages. Although our molecular work support a direct regulation HCN1 by *Nr2f1*, it is complex to precisely dissect changes in membrane-ion channels, priming the start of abnormal network excitability, from those that are concomitantly changing by intrinsic or homeostatic plasticity. Nevertheless, we would like to propose that decreased HCN1 levels could act as initiator of the deregulation in network as well as intrinsic activity observed in mutant cortices. This would imply that the alterations in expression levels we observed in other voltage-gated ion channels are the product of additional network-activity-driven compensatory mechanisms occurring after P1 or P4. In other words, we hypothesize that changes in ion-channel expression of pyramidal glutamatergic neurons at P5–P8 are due to homeostatic and/or intrinsic plasticity changes following earlier alterations in overall network activity state. For example, experimental evidence from the developing hippocampus suggests that the induction of increased bursting activity and seizures alters persistently HCN channel expression and voltage-dependence of I_h_ current (Arnold *et al*., 2019; Chen *et al*., 2001). Acting as a direct transcriptional regulator of HCN1, we propose that Nr2f1 together with other transcriptional regulators, such as Tbr1 (Fazel Darbandi et al., 2018), could influence spontaneous network activity intrinsic to the somatosensory cortex by contributing to the establishment of the bioelectrical state of glutamatergic neurons at the time point, and probably before, the start of cortical network formation. Moreover, intrinsic pacemaker properties through HCN1 expression has been hypothesized to generate and drive distinct synchronized activity patterns in immature networks (Luhmann *et al*., 2016). Although we identified HCN1 as a direct effector of the intrinsic electrophysiological phenotype observed in Nr2f1-deficient cortices, an additional complement of ion channels, regulated at the transcriptional or posttranscriptional level, might actively contribute to the impairment of network activity.

Importantly, genetic or pharmacological dysfunction in HCN channels in mice results in behavioral phenotypes associated to somatosensory-motor coordination, e.g. reduced forelimb reaching accuracy and atypical movements during a single-pellet skill reaching task (Boychuk *et al*., 2017). This is very analogous to the behavioral phenotypes displayed by *Nr2f1cKO* mice (Tomassy *et al*., 2010) and symptoms of NR2F1 haploinsufficient patients (Bosch *et al*., 2014; Chen *et al*., 2016), suggesting that Nr2f1 might influence cortical maps involving voluntary movements by allowing coordinated skilled behavior *via* the action of HCN1.

Together with the cellular and molecular role of Nr2f1 during intrinsic area patterning, this study has unraveled a key and novel functional role of this transcriptional regulator in cortical intrinsic excitability and spontaneous activity that might affect axonal connectivity and circuit formation necessary for behavioral abnormalities described in mice (Contesse *et al*., 2019; Flore *et al*., 2017; Tomassy *et al*., 2010) and, possibly, in humans (Chen *et al*., 2016; Bosch *et al*., 2016).

## ACKNOWLEDGMENTS

We thank Dr. Marteen Kole and Dr. Nathalie Dehorter for critical reading of the manuscript and members of the Studer and Frick laboratories for stimulating discussions. We also thank Guillermina Lopez-Bendito for insightful suggestions. We are grateful to the iBV PRISM Microscopy facility and the Bordeaux Imaging Center (BIC), supported by Labex-BRAIN (ANR-10-LABEX-43), for the acquisition of some imaging data. The authors also thank the animal- and genotyping facilities of the Neurocentre Magendie (supported by INSERM and LabEX-BRAIN ANR-10-LABX-43) and of the iBV. This work was funded by an ANR (Agence National de la Recherche) grant # ANR-13-BSV4-0011-01 to M.S. and X.L, by the “Fondation Recherche Médicale; Equipe FRM 2015 grant #DEQ20150331750 to M.S.; by an ‘‘Investments for the Future’’ LabEx SIGNALIFE (grant ANR-11-LABX-0028-01) to M.S., by the “Initiative d’Excellence (IdEx) de ĺUniversité de Bordeaux” fellowship to I.D.P., and by the Inserm endowment to A.F. Address of the corresponding author: (iBV - Institut de Biologie Valrose, Univ. Nice Sophia Antipolis, Centre de Biochimie; UFR Sciences, Parc Valrose, 28 avenue Valrose, 06108 Nice Cedex 2, France).

## Author Contributions

Conceptualization, Methodology, Validation, & Formal Analysis, I.D.P., C.T., E.M., A.M., C.A., M.S., X.L, A.F.; Investigation, I.D.P., C.T., E.M., A.M., C.F., G.T.; Resources, M.S., A.F.; Writing – Original Draft & Editing, I.D.P., C.T., E.M., A.M., M.S.; Writing – Review, X.L. and A.F.; Visualization, I.D.P., C.T., E.M., A.M.; Funding Acquisition, M.S., X.L., A.F, I.D.P.

## Declaration of Interests

The corresponding author declares non-financial competing interest on behalf of all authors.

## Notes

#### Summary of Updates

Major revision with changes in the overall logical organization of the data and manuscript. More experimental and statistical details, but no additional experiments.

## REFERENCES

Alfano, C., Magrinelli, E., Harb, K., Hevner, R. F. and Studer, M. (2014) ‘Postmitotic control of sensory area specification during neocortical development’, Nat Commun, 5, pp. 5632.

Alfano, C. and Studer, M. (2012) ‘Neocortical arealization: Evolution, mechanisms and open questions’, Dev Neurobiol.

Alfano, C., Viola, L., Heng, J. I., Pirozzi, M., Clarkson, M., Flore, G., De Maio, A., Schedl, A., Guillemot, F. and Studer, M. (2011) ‘COUP-TFI promotes radial migration and proper morphology of callosal projection neurons by repressing Rnd2 expression’, Development, 138(21), pp. 4685–97.

Allene, C., Cattani, A., Ackman, J. B., Bonifazi, P., Aniksztejn, L., Ben-Ari, Y. and Cossart, R. (2008) ‘Sequential generation of two distinct synapse-driven network patterns in developing neocortex’, J Neurosci, 28(48), pp. 12851–63.

Allene, C. and Cossart, R. (2010) ‘Early NMDA receptor-driven waves of activity in the developing neocortex: physiological or pathological network oscillations?’, J Physiol, 588(Pt 1), pp. 83–91.

Anderson, O. D. 1975. Moving Average Processes. The Statistician.

Andreae, L. C. and Burrone, J. (2018) ‘The role of spontaneous neurotransmission in synapse and circuit development’, J Neurosci Res, 96(3), pp. 354–359.

Anton-Bolanos, N., Sempere-Ferrandez, A., Guillamon-Vivancos, T., Martini, F. J., Perez-Saiz, L., Gezelius, H., Filipchuk, A., Valdeolmillos, M. and Lopez-Bendito, G. (2019) ‘Prenatal activity from thalamic neurons governs the emergence of functional cortical maps in mice’, Science, 364(6444), pp. 987–990.

Arlotta, P., Molyneaux, B. J., Chen, J., Inoue, J., Kominami, R. and Macklis, J. D. (2005) ‘Neuronal subtype-specific genes that control corticospinal motor neuron development in vivo’, Neuron, 45(2), pp. 207–21.

Armentano, M., Chou, S. J., Tomassy, G. S., Leingartner, A., O’Leary, D. D. and Studer, M. (2007) ‘COUP-TFI regulates the balance of cortical patterning between frontal/motor and sensory areas’, Nat Neurosci, 10(10), pp. 1277–86.

Arnold, E. C., McMurray, C., Gray, R. and Johnston, D. (2019) ‘Epilepsy-Induced Reduction in HCN Channel Expression Contributes to an Increased Excitability in Dorsal, But Not Ventral, Hippocampal CA1 Neurons’, eNeuro, 6(2).

Bender, R. A. and Baram, T. Z. (2008) ‘Hyperpolarization activated cyclic-nucleotide gated (HCN) channels in developing neuronal networks’, Prog Neurobiol, 86(3), pp. 129–40.

Berger, T., Larkum, M. E. and Luscher, H. R. (2001) ‘High I(h) channel density in the distal apical dendrite of layer V pyramidal cells increases bidirectional attenuation of EPSPs’, J Neurophysiol, 85(2), pp. 855–68.

Blankenship, A. G., Hamby, A. M., Firl, A., Vyas, S., Maxeiner, S., Willecke, K. and Feller, M. B. (2011) ‘The role of neuronal connexins 36 and 45 in shaping spontaneous firing patterns in the developing retina’, J Neurosci, 31(27), pp. 9998–10008.

Boillot, M., Lee, C. Y., Allene, C., Leguern, E., Baulac, S. and Rouach, N. (2016) ‘LGI1 acts presynaptically to regulate excitatory synaptic transmission during early postnatal development’, Sci Rep, 6, pp. 21769.

Bosch, D. G., Boonstra, F. N., de Leeuw, N., Pfundt, R., Nillesen, W. M., de Ligt, J., Gilissen, C., Jhangiani, S., Lupski, J. R., Cremers, F. P. and de Vries, B. B. (2016) ‘Novel genetic causes for cerebral visual impairment’, Eur J Hum Genet, 24(5), pp. 660–5.

Bosch, D. G., Boonstra, F. N., Gonzaga-Jauregui, C., Xu, M., de Ligt, J., Jhangiani, S., Wiszniewski, W., Muzny, D. M., Yntema, H. G., Pfundt, R., Vissers, L. E., Spruijt, L., Blokland, E. A., Chen, C. A., Baylor-Hopkins Center for Mendelian, G., Lewis, R. A., Tsai, S. Y., Gibbs, R. A., Tsai, M. J., Lupski, J. R., Zoghbi, H. Y., Cremers, F. P., de Vries, B. B. and Schaaf, C. P. (2014) ‘NR2F1 mutations cause optic atrophy with intellectual disability’, Am J Hum Genet, 94(2), pp. 303–9.

Boychuk, J. A., Farrell, J. S., Palmer, L. A., Singleton, A. C., Pittman, Q. J. and Teskey, G. C. (2017) ‘HCN channels segregate stimulation-evoked movement responses in neocortex and allow for coordinated forelimb movements in rodents’, J Physiol, 595(1), pp. 247–263.

Brager, D. H., Akhavan, A. R. and Johnston, D. (2012) ‘Impaired dendritic expression and plasticity of h-channels in the fmr1(-/y) mouse model of fragile X syndrome’, Cell Rep, 1(3), pp. 225–33.

Brager, D. H. and Johnston, D. (2007) ‘Plasticity of intrinsic excitability during long-term depression is mediated through mGluR-dependent changes in I(h) in hippocampal CA1 pyramidal neurons’, J Neurosci, 27(51), pp. 13926–37.

Chen, C. A., Bosch, D. G., Cho, M. T., Rosenfeld, J. A., Shinawi, M., Lewis, R. A., Mann, J., Jayakar, P., Payne, K., Walsh, L., Moss, T., Schreiber, A., Schoonveld, C., Monaghan, K. G., Elmslie, F., Douglas, G., Boonstra, F. N., Millan, F., Cremers, F. P., McKnight, D., Richard, G., Juusola, J., Kendall, F., Ramsey, K., Anyane-Yeboa, K., Malkin, E., Chung, W. K., Niyazov, D., Pascual, J. M., Walkiewicz, M., Veluchamy, V., Li, C., Hisama, F. M., de Vries, B. B. and Schaaf, C. (2016) ‘The expanding clinical phenotype of Bosch-Boonstra-Schaaf optic atrophy syndrome: 20 new cases and possible genotype-phenotype correlations’, Genet Med.

Chen, K., Aradi, I., Thon, N., Eghbal-Ahmadi, M., Baram, T. Z. and Soltesz, I. (2001) ‘Persistently modified h-channels after complex febrile seizures convert the seizure-induced enhancement of inhibition to hyperexcitability’, Nat Med, 7(3), pp. 331–7.

Cho, M. W. and Choi, M. Y. (2012) ‘Spontaneous organization of the cortical structure through endogenous neural firing and gap junction transmission’, Neural Netw, 31, pp. 46–52.

Christophe, E., Doerflinger, N., Lavery, D. J., Molnar, Z., Charpak, S. and Audinat, E. (2005) ‘Two populations of layer v pyramidal cells of the mouse neocortex: development and sensitivity to anesthetics’, J Neurophysiol, 94(5), pp. 3357–67.

Computing, R. F. f. S. 2018. R: A Language and Environment for Statistical Computing. R Foundation for Statistical Computing.

Contesse, T., Ayrault, M., Mantegazza, M., Studer, M. and Deschaux, O. (2019) ‘Hyperactive and anxiolytic-like behaviors result from loss of COUP-TFI/Nr2f1 in the mouse cortex’, Genes Brain Behav, pp. e12556.

Corlew, R., Bosma, M. M. and Moody, W. J. (2004) ‘Spontaneous, synchronous electrical activity in neonatal mouse cortical neurones’, J Physiol, 560(Pt 2), pp. 377–90.

Del Pino, I., Brotons-Mas, J. R., Marques-Smith, A., Marighetto, A., Frick, A., Marin, O. and Rico, B. (2017) ‘Abnormal wiring of CCK(+) basket cells disrupts spatial information coding’, Nat Neurosci, 20(6), pp. 784–792.

Del Pino, I., Garcia-Frigola, C., Dehorter, N., Brotons-Mas, J. R., Alvarez-Salvado, E., Martinez de Lagran, M., Ciceri, G., Gabaldon, M. V., Moratal, D., Dierssen, M., Canals, S., Marin, O. and Rico, B. (2013) ‘Erbb4 deletion from fast-spiking interneurons causes schizophrenia-like phenotypes’, Neuron, 79(6), pp. 1152–68.

Dumitrescu, A. S., Evans, M. D. and Grubb, M. S. (2016) ‘Evaluating Tools for Live Imaging of Structural Plasticity at the Axon Initial Segment’, Front Cell Neurosci, 10, pp. 268.

Faedo, A., Tomassy, G. S., Ruan, Y., Teichmann, H., Krauss, S., Pleasure, S. J., Tsai, S. Y., Tsai, M. J., Studer, M. and Rubenstein, J. L. (2008) ‘COUP-TFI coordinates cortical patterning, neurogenesis, and laminar fate and modulates MAPK/ERK, AKT, and beta-catenin signaling’, Cereb Cortex, 18(9), pp. 2117–31.

Fan, J., Stemkowski, P. L., Gandini, M. A., Black, S. A., Zhang, Z., Souza, I. A., Chen, L. and Zamponi, G. W. (2016) ‘Reduced Hyperpolarization-Activated Current Contributes to Enhanced Intrinsic Excitability in Cultured Hippocampal Neurons from PrP(-/-) Mice’, Front Cell Neurosci, 10, pp. 74.

Fan, Y., Fricker, D., Brager, D. H., Chen, X., Lu, H. C., Chitwood, R. A. and Johnston, D. (2005) ‘Activity-dependent decrease of excitability in rat hippocampal neurons through increases in I(h)’, Nat Neurosci, 8(11), pp. 1542–51.

Favuzzi, E., Deogracias, R., Marques-Smith, A., Maeso, P., Jezequel, J., Exposito-Alonso, D., Balia, M., Kroon, T., Hinojosa, A. J., E, F. M. and Rico, B. (2019) ‘Distinct molecular programs regulate synapse specificity in cortical inhibitory circuits’, Science, 363(6425), pp. 413–417.

Fazel Darbandi, S., Robinson Schwartz, S. E., Qi, Q., Catta-Preta, R., Pai, E. L., Mandell, J. D., Everitt, A., Rubin, A., Krasnoff, R. A., Katzman, S., Tastad, D., Nord, A. S., Willsey, A. J., Chen, B., State, M. W., Sohal, V. S. and Rubenstein, J. L. R. (2018) ‘Neonatal Tbr1 Dosage Controls Cortical Layer 6 Connectivity’, Neuron, 100(4), pp. 831–845.e7.

Flore, G., Di Ruberto, G., Parisot, J., Sannino, S., Russo, F., Illingworth, E. A., Studer, M. and De Leonibus, E. (2017) ‘Gradient COUP-TFI Expression Is Required for Functional Organization of the Hippocampal Septo-Temporal Longitudinal Axis’, Cereb Cortex, 27(2), pp. 1629–1643.

Gavello, D., Calorio, C., Franchino, C., Cesano, F., Carabelli, V., Carbone, E. and Marcantoni, A. (2018) ‘Early Alterations of Hippocampal Neuronal Firing Induced by Abeta42’, Cereb Cortex, 28(2), pp. 433–446.

Gavello, D., Rojo-Ruiz, J., Marcantoni, A., Franchino, C., Carbone, E. and Carabelli, V. (2012) ‘Leptin counteracts the hypoxia-induced inhibition of spontaneously firing hippocampal neurons: a microelectrode array study’, PLoS One, 7, pp. e41530.

Greig, L. C., Woodworth, M. B., Galazo, M. J., Padmanabhan, H. and Macklis, J. D. (2013) ‘Molecular logic of neocortical projection neuron specification, development and diversity’, Nat Rev Neurosci, 14(11), pp. 755–69.

Grubb, M. S. and Burrone, J. (2010) ‘Activity-dependent relocation of the axon initial segment fine-tunes neuronal excitability’, Nature, 465(7301), pp. 1070–4.

Gulledge, A. T. and Bravo, J. J. (2016) ‘Neuron Morphology Influences Axon Initial Segment Plasticity’, eNeuro, 3(1).

Hamada, M. S., Goethals, S., de Vries, S. I., Brette, R. and Kole, M. H. (2016) ‘Covariation of axon initial segment location and dendritic tree normalizes the somatic action potential’, Proc Natl Acad Sci U S A, 113(51), pp. 14841–14846.

Harb, K., Magrinelli, E., Nicolas, C. S., Lukianets, N., Frangeul, L., Pietri, M., Sun, T., Sandoz, G., Grammont, F., Jabaudon, D., Studer, M. and Alfano, C. (2016) ‘Area-specific development of distinct projection neuron subclasses is regulated by postnatal epigenetic modifications’, Elife, 5, pp. e09531.

Jabaudon, D. (2017) ‘Fate and freedom in developing neocortical circuits’, Nat Commun, 8, pp. 16042.

Kasper, E. M., Larkman, A. U., Lubke, J. and Blakemore, C. (1994) ‘Pyramidal neurons in layer 5 of the rat visual cortex. II. Development of electrophysiological properties’, J Comp Neurol, 339(4), pp. 475–94.

Kirischuk, S., Sinning, A., Blanquie, O., Yang, J. W., Luhmann, H. J. and Kilb, W. (2017) ‘Modulation of Neocortical Development by Early Neuronal Activity: Physiology and Pathophysiology’, Front Cell Neurosci, 11, pp. 379.

Kirkby, L. A., Sack, G. S., Firl, A. and Feller, M. B. (2013) ‘A role for correlated spontaneous activity in the assembly of neural circuits’, Neuron, 80(5), pp. 1129–44.

Kole, M. H. and Stuart, G. J. (2012) ‘Signal processing in the axon initial segment’, Neuron, 73(2), pp. 235–47.

Kordeli, E., Lambert, S. and Bennett, V. (1995) ‘AnkyrinG. A new ankyrin gene with neural-specific isoforms localized at the axonal initial segment and node of Ranvier’, J Biol Chem, 270(5), pp. 2352–9.

Kuba, H. (2010) ‘Plasticity at the axon initial segment’, Commun Integr Biol, 3(6), pp. 597–8.

Kuba, H., Oichi, Y. and Ohmori, H. (2010) ‘Presynaptic activity regulates Na(+) channel distribution at the axon initial segment’, Nature, 465(7301), pp. 1075–8.

Li, H., Fertuzinhos, S., Mohns, E., Hnasko, T. S., Verhage, M., Edwards, R., Sestan, N. and Crair, M. B. (2013) ‘Laminar and columnar development of barrel cortex relies on thalamocortical neurotransmission’, Neuron, 79(5), pp. 970–86.

Loots, G. and Ovcharenko, I. (2007) ‘ECRbase: database of evolutionary conserved regions, promoters, and transcription factor binding sites in vertebrate genomes’, Bioinformatics, 23(1), pp. 122–4.

Luhmann, H. J. and Khazipov, R. (2018) ‘Neuronal activity patterns in the developing barrel cortex’, Neuroscience, 368, pp. 256–267.

Luhmann, H. J., Sinning, A., Yang, J. W., Reyes-Puerta, V., Stuttgen, M. C., Kirischuk, S. and Kilb, W. (2016) ‘Spontaneous Neuronal Activity in Developing Neocortical Networks: From Single Cells to Large-Scale Interactions’, Front Neural Circuits, 10, pp. 40.

Magee, J. C. (1998) ‘Dendritic hyperpolarization-activated currents modify the integrative properties of hippocampal CA1 pyramidal neurons’, J Neurosci, 18(19), pp. 7613–24.

Montemayor, C., Montemayor, O. A., Ridgeway, A., Lin, F., Wheeler, D. A., Pletcher, S. D. and Pereira, F. A. (2010) ‘Genome-wide analysis of binding sites and direct target genes of the orphan nuclear receptor NR2F1/COUP-TFI’, PLoS One, 5(1), pp. e8910.

Moody, W. J. and Bosma, M. M. (2005) ‘Ion channel development, spontaneous activity, and activity-dependent development in nerve and muscle cells’, Physiol Rev, 85(3), pp. 883–941.

Murase, S., Lantz, C. L., Kim, E., Gupta, N., Higgins, R., Stopfer, M., Hoffman, D. A. and Quinlan, E. M. (2016) ‘Matrix Metalloproteinase-9 Regulates Neuronal Circuit Development and Excitability’, Mol Neurobiol, 53(5), pp. 3477–3493.

O’Leary, D. D. and Sahara, S. (2008) ‘Genetic regulation of arealization of the neocortex’, Curr Opin Neurobiol, 18(1), pp. 90–100.

Ovcharenko, I., Nobrega, M. A., Loots, G. G. and Stubbs, L. (2004) ‘ECR Browser: a tool for visualizing and accessing data from comparisons of multiple vertebrate genomes’, Nucleic Acids Res, 32(Web Server issue), pp. W280–6.

Parisot, J., Flore, G., Bertacchi, M. and Studer, M. (2017) ‘COUP-TFI mitotically regulates production and migration of dentate granule cells and modulates hippocampal Cxcr4 expression’, Development, 144(11), pp. 2045–2058.

Pearson, R. A., Catsicas, M., Becker, D. L., Bayley, P., Luneborg, N. L. and Mobbs, P. (2004) ‘Ca(2+) signalling and gap junction coupling within and between pigment epithelium and neural retina in the developing chick’, Eur J Neurosci, 19(9), pp. 2435–45.

Poolos, N. P., Bullis, J. B. and Roth, M. K. (2006) ‘Modulation of h-channels in hippocampal pyramidal neurons by p38 mitogen-activated protein kinase’, J Neurosci, 26(30), pp. 7995–8003.

Poolos, N. P., Migliore, M. and Johnston, D. (2002) ‘Pharmacological upregulation of h-channels reduces the excitability of pyramidal neuron dendrites’, Nat Neurosci, 5(8), pp. 767–74.

Quandt, K., Frech, K., Karas, H., Wingender, E. and Werner, T. (1995) ‘MatInd and MatInspector: new fast and versatile tools for detection of consensus matches in nucleotide sequence data’, Nucleic Acids Res, 23(23), pp. 4878–84.

Roerig, B. and Feller, M. B. (2000) ‘Neurotransmitters and gap junctions in developing neural circuits’, Brain Res Brain Res Rev, 32(1), pp. 86–114.

Schindelin, J., Arganda-Carreras, I., Frise, E., Kaynig, V., Longair, M., Pietzsch, T., Preibisch, S., Rueden, C., Saalfeld, S., Schmid, B., Tinevez, J. Y., White, D. J., Hartenstein, V., Eliceiri, K., Tomancak, P. and Cardona, A. (2012) ‘Fiji: an open-source platform for biological-image analysis’, Nat Methods, 9(7), pp. 676–82.

Schneggenburger, R. and Rosenmund, C. (2015) ‘Molecular mechanisms governing Ca(2+) regulation of evoked and spontaneous release’, Nat Neurosci, 18(7), pp. 935–41.

Shah, M. M. (2014) ‘Cortical HCN channels: function, trafficking and plasticity’, J Physiol, 592(13), pp. 2711–9.

Simi, A. and Studer, M. (2018) ‘Developmental genetic programs and activity-dependent mechanisms instruct neocortical area mapping’, Curr Opin Neurobiol, 53, pp. 96–102.

Tomassy, G. S., De Leonibus, E., Jabaudon, D., Lodato, S., Alfano, C., Mele, A., Macklis, J. D. and Studer, M. (2010) ‘Area-specific temporal control of corticospinal motor neuron differentiation by COUP-TFI’, Proc Natl Acad Sci U S A, 107(8), pp. 3576–81.

Tritsch, N. X., Yi, E., Gale, J. E., Glowatzki, E. and Bergles, D. E. (2007) ‘The origin of spontaneous activity in the developing auditory system’, Nature, 450(7166), pp. 50–5.

Uhlen, P., Fritz, N., Smedler, E., Malmersjo, S. and Kanatani, S. (2015) ‘Calcium signaling in neocortical development’, Dev Neurobiol, 75(4), pp. 360–8.

van Welie, I., Remme, M. W., van Hooft, J. A. and Wadman, W. J. (2006) ‘Different levels of Ih determine distinct temporal integration in bursting and regular-spiking neurons in rat subiculum’, J Physiol, 576(Pt 1), pp. 203–14.

Wang, Z., Xu, N. L., Wu, C. P., Duan, S. and Poo, M. M. (2003) ‘Bidirectional changes in spatial dendritic integration accompanying long-term synaptic modifications’, Neuron, 37(3), pp. 463–72

Wefelmeyer, W., Puhl, C. J. and Burrone, J. (2016) ‘Homeostatic Plasticity of Subcellular Neuronal Structures: From Inputs to Outputs’, Trends Neurosci, 39(10), pp. 656–667.

Williams, S. R. and Stuart, G. J. (2000) ‘Site independence of EPSP time course is mediated by dendritic I(h) in neocortical pyramidal neurons’, J Neurophysiol, 83(5), pp. 3177–82.

Williams, S. R. and Stuart, G. J. (2003) ‘Voltage- and site-dependent control of the somatic impact of dendritic IPSPs’, J Neurosci, 23(19), pp. 7358–67.

Yamada, R. and Kuba, H. (2016) ‘Structural and Functional Plasticity at the Axon Initial Segment’, Front Cell Neurosci, 10, pp. 250.

Yamamoto, N. and Lopez-Bendito, G. (2012) ‘Shaping brain connections through spontaneous neural activity’, Eur J Neurosci, 35(10), pp. 1595–604.

Zhou, D., Lambert, S., Malen, P. L., Carpenter, S., Boland, L. M. and Bennett, V. (1998) ‘AnkyrinG is required for clustering of voltage-gated Na channels at axon initial segments and for normal action potential firing’, J Cell Biol, 143(5), pp. 1295–304.

