## Supplementary Figures and Tables for "COUP-TFI/Nr2f1 orchestrates intrinsic neuronal activity during cortical area patterning"

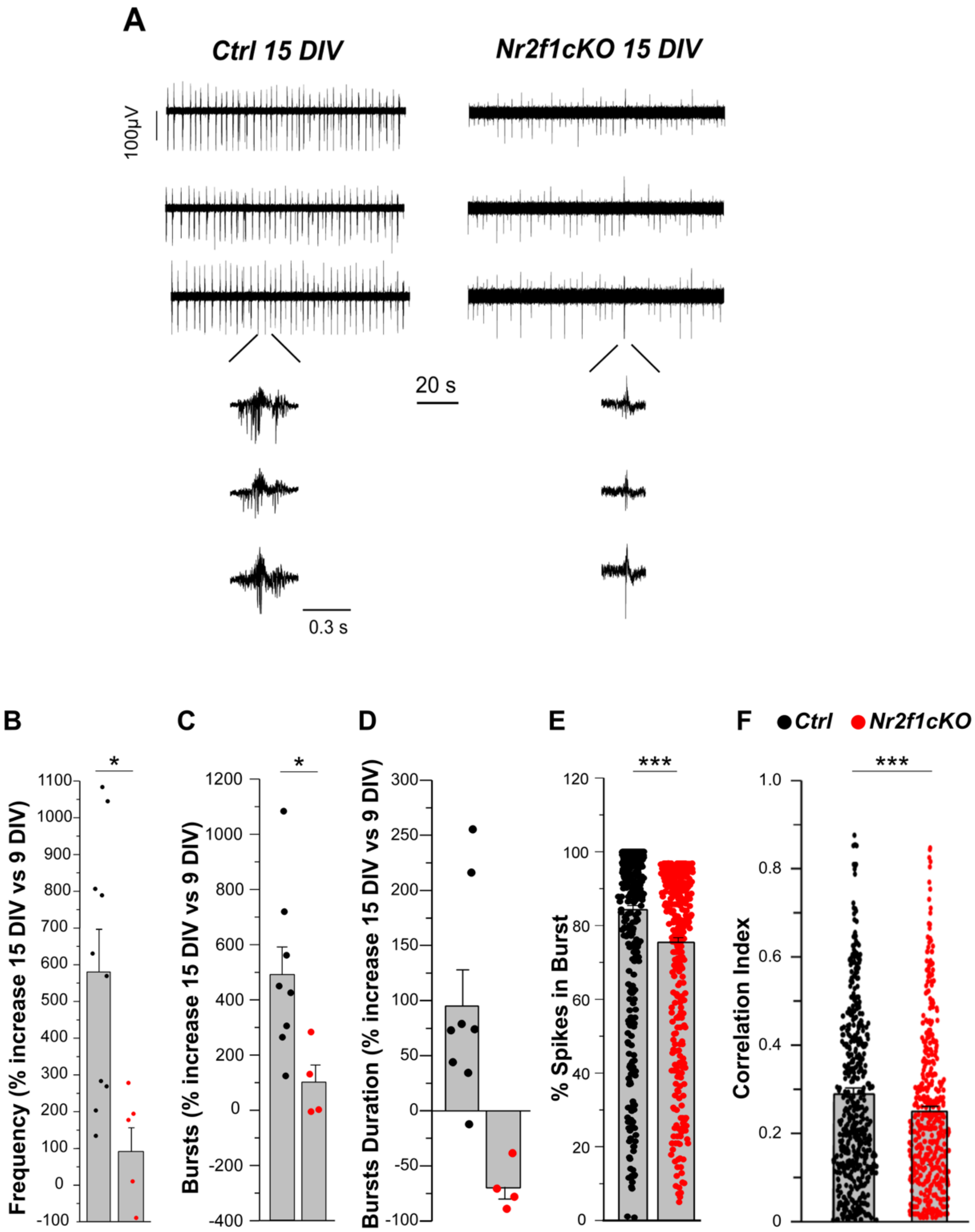

***Figure S1. Impaired intrinsic network and bursting activity recorded in DIV15 cultured Nr2f1-deficient cortical neurons.*** **A**) Spontaneous firing of *Ctrl* (left) and *Nr2f1cKO* (right) cortical network recorded by three representative MEA electrodes a 15DIV. In insets, representative bursts shown at expanded scale. **B-F**) Firing parameters measured in *Ctrl* and *Nr2f1cKO* cortical networks are compared: frequency (% increase 15 DIV vs 9 DIV) (*Ctrl*, 580.155 ±116.167, $N_{MEA}$ = 10; *Nr2f1cKO*, 91.455 ± 64.271, $N_{MEA}$ =6), bursts (% increase 15 DIV vs 9 DIV) (*Ctrl*, 489.058 ±100.530, $N_{MEA}$ = 8; *Nr2f1cKO*, 100.682 ± 60.654, $N_{MEA}$ =4), burst duration (% increase 15 DIV vs 9 DIV) (*Ctrl*, 95.245 ±32.591, $N_{MEA}$ =8; *Nr2f1cKO*, -69.773 ± 10.813, $N_{MEA}$ =4), % spikes in burst (*Ctrl*, 84.292 ±1.137, $N_{channels}$ = 458, from $N_{MEA}$ = 10; *Nr2f1cKO*, 78.711 ± 1.188, $N_{channels}$ =447, from $N_{MEA}$ = 6), cross correlation index (*Ctrl*, 0.289 ±0.010, $N_{channels}$ = 727, from $N_{MEA}$ = 10; *Nr2f1cKO*, 0.250 ± 0.009, $N_{channels}$ =626, from $N_{MEA}$ = 6) (* P<0.05; ***P<0.001, unpaired T-test). Data are represented as mean ± SEM. ***See also Figure 1.***

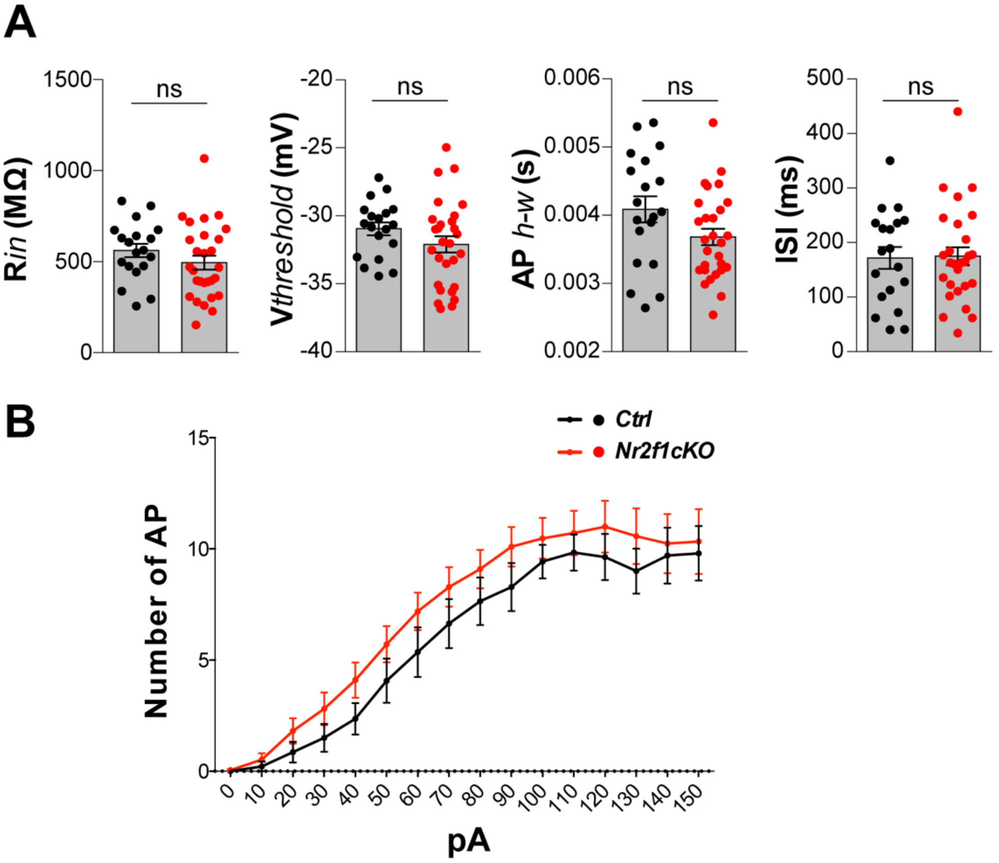

***Figure S2. Specific intrinsic excitability features and input-output relationship remain unaltered in Nr2f1 deficient layer V pyramidal neurons (LVPNs).*** **A**) LVPNs in *Ctrl* (in black) and *Nr2f1cKO* (in red) display normal input resistance and threshold for action potential and inter-spike interval. We observed a tendency of *Nr2f1cKO* LVPNs to display a reduced half-width of action potential when compared to *Ctrl* LVPNs (*P = 0.066*) and a significant decreased AP*amp* (**P = 0.04441*) measured from resting membrane potential (N = 19 *Ctrl*, N = 28 *Nr2f1cKO*, two-tailed unpaired T-test or U Mann-Whitney test). **B**) Input/output function is not significantly different between *Nr2f1cKO* LVPNs and *Ctrl* LVPNs (N = 19 *Ctrl*, N = 28 *Nr2f1cKO*, two-way ANOVA). Data are represented as mean ± SEM. R*in*, input resistance; V*threshold*, threshold for action potential; AP*h-w*, half-width of action potential; ISI, inter-spike interval. ***See also Figure 3 and TableS1.***

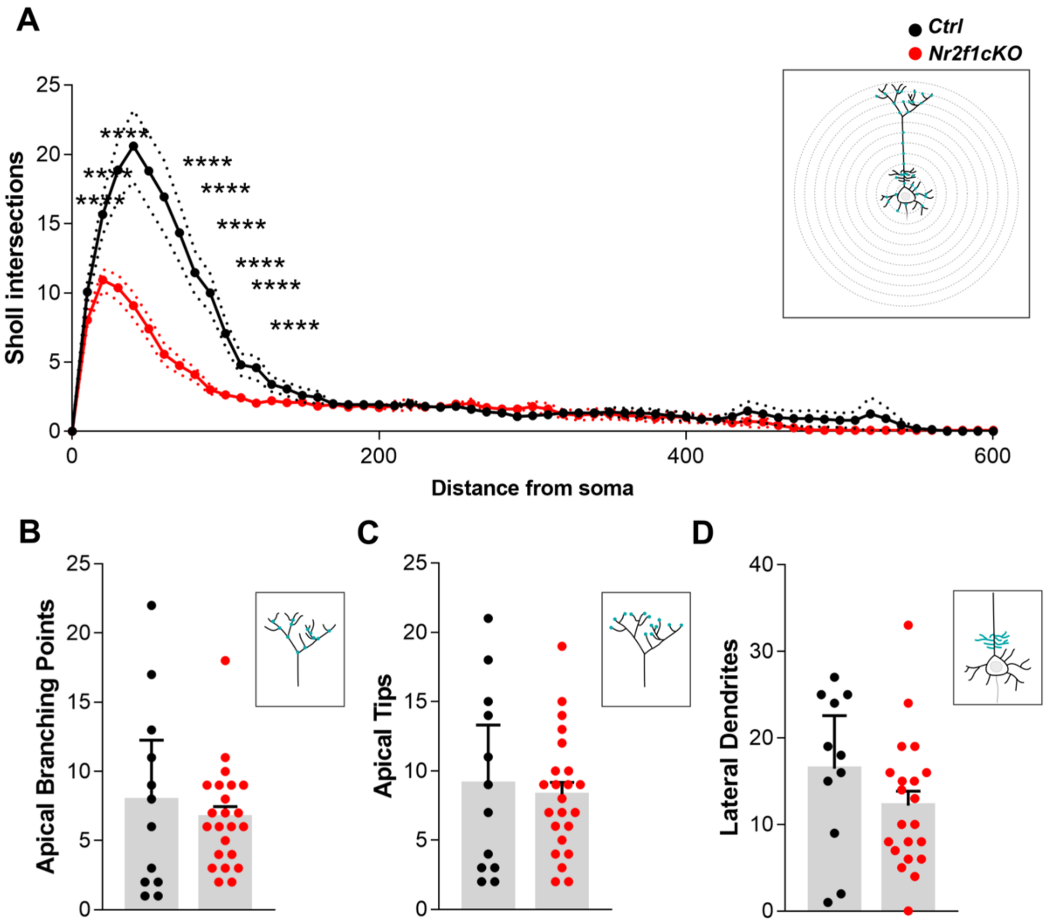

***Figure S3. Apical dendritic complexity of layer V pyramidal neurons (LVPNs) is not affected in Nr2f1 mutant brains.*** **A**) Sholl analysis shows no differences in complexity between *Ctrl* and *Nr2f1cKO* LVPNs, relatively to distal dendrites. (ns *P > 0.5*) **B-D**) Number of Apical Branching Points (*Ctrl,* 7.92 ± 1.97**;** *Nr2f1cKO* 6.7 ± 0.75**;** P = 0.57), Apical Terminal Tips (*Ctrl,* 9.08 ± 1.93**,** *Nr2f1cKO* 8.26 ± 0.87**;** P = 0.70) and Lateral dendrites (*Ctrl,* 16.45 ± 2.75**;** *Nr2f1cKO,* 12.19 ± 1.65; *P =0.20,*) are unaffected by the loss of *Nr2f1* (N = 12 *Ctrl*, N = 23 *Nr2f1cKO;* N defined as number of analyzed cells). P-values are calculated by 2way ANOVA test (**A**) or Welch's unequal variances t-test (**B-D**). Data are represented as mean ± SEM. ***See also Figure 4 and Table S2..***

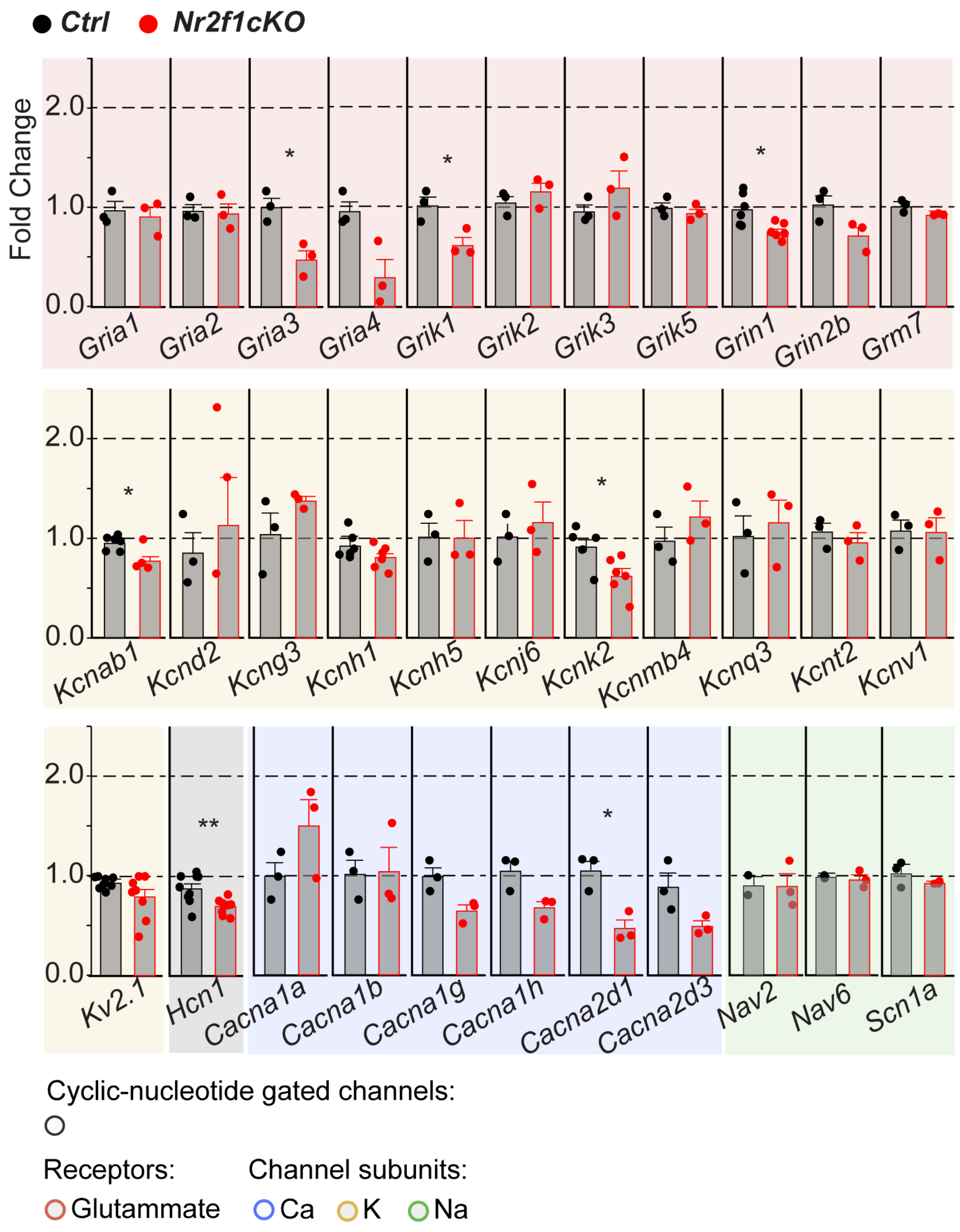

***Figure S4. qPCR quantification of cortical ion channels in Nr2f1 mutant cortices.*** Single qPCR quantification of P7 *Ctrl* vs *Nr2f1cKO* expression of chosen cortically (deep layers) expressed ion channel subunits and Glutamate receptors. *Ctrl* mean value is normalized to 1 to calculate fold changes in *Nr2f1cKO* conditions. N≥3. Statistical significance is tested doing paired t-test with P-value adjusted according to Benjamini–Hochberg procedure. *- P-Value < 0.05. P – Postnatal day. Data are represented as mean ± SEM. ***See also Figure 6.***

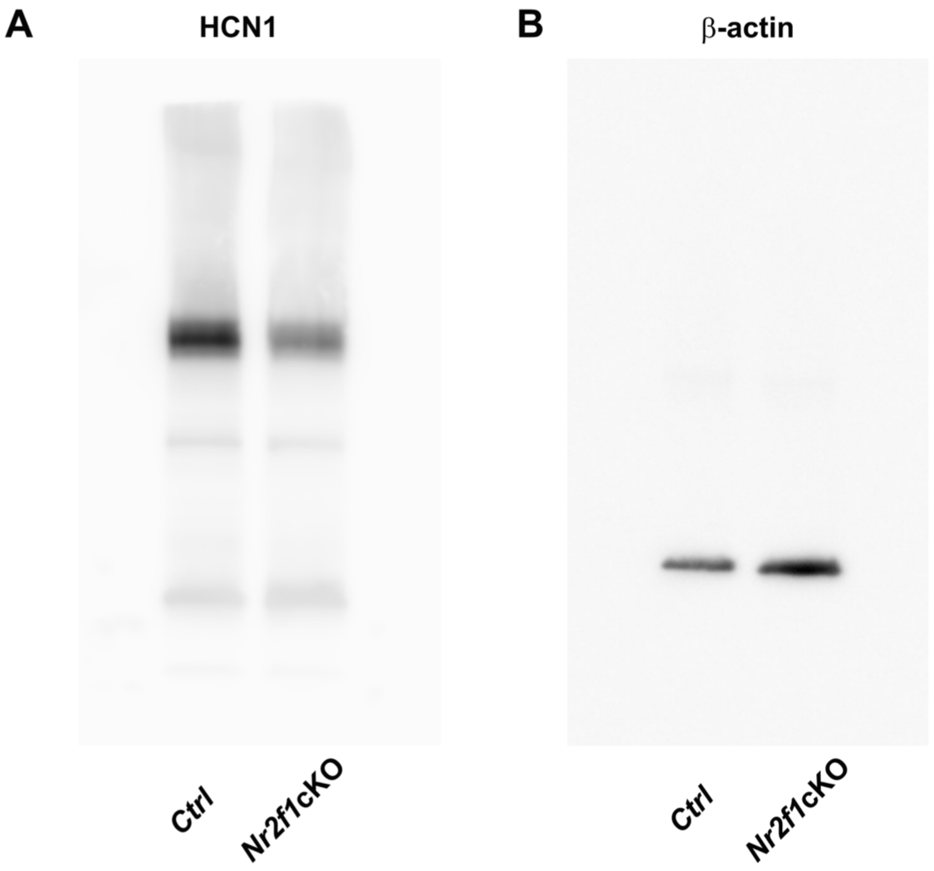

***Figure S5. Full lane western blot of cortical ion channel HCN1.*** **A**) Western Blot of HCN1 *in Ctrl and Nr2f1 mutant cortices.* Note specific downregulationof HCN1 in the mutant brain. B) Western Blot of b-actin *in Ctrl and Nr2f1 mutant cortices* as an internal control. ***See also Figure 6.***

| | ***Ctrl*** | ***cKO*** | p-value | test |
|---|---|---|---|---|
| $V_{rest}$ (mV) | -60.26 ± 1.18 | -54.62 ± 1.13 | 0.0013 (*) | t-test |
| $R_{in}$ (MΩ) | 561.2 ± 36.66 | 494.80 ± 38.64 | 0.2410 | t-test |
| Latency (ms) | 133.5 ± 15.46 | 145.9 ± 23.59 | 0.9023 | Mann-Whitney |
| Rheobase (pA) | 61.05 ± 9.08 | 29.64 ± 3.47 | 0.001(***) | t-test |
| $V_{threshold}$ (mV) | -30.95 ± 0.48 | -32.09 ± 0.60 | 0.1750 | t-test |
| $AP_{amp}$ (mV) | 66.65 ± 1.62 | 69.1 ± 1.11 | 0.2042 | t-test |
| $AP_{\text{half-width}}$ (ms) | 0.0041±0.0002 | 0.0037±0.0001 | 0.0659 | t-test |
| Inter-spike-interval(ISI) (ms) | 172.0 ± 19.92 | 175.1 ± 16.42 | 0.9529 | Mann-Whitney |
| Sag (mV) | 8.365 ± 0.89 | 5.85 ± 0.90 | 0.0418(*) | Mann-Whitney |
| $\tau_m$ (ms) | 28.42 ± 3.26 | 31.14 ± 3.70 | 0.6193 | Mann-Whitney |

***Table S1*. *Nr2f1* deficiency leads to specific changes in intrinsic electrophysiological properties in pyramidal neurons within the immature somatosensory cortex.** Intrinsic electrophysiological properties of layer V pyramidal neurons in acute brain slices of the immature somatosensory cortex from *Ctrl* (19 cells from 8 different brains) and *Nr2f1cKO* (28 cells from 7 different brains) P5-P8 mouse pups. $V_{rest}$: resting membrane potential; $R_{in}$ : input resistance; $V_{threshold}$ : action potential threshold; $AP_{amp}$ : action potential amplitude; $AP_{\text{half-width}}$: action potential half-width; $\tau_m$: membrane time constant. ***See also Figure 3 and S2.***

| *Sholl analysis statistics: 0-120 mm* | | | | | | |
|---|---|---|---|---|---|---|
| | ***Ctrl mean values*** | ***Nr2f1 cKO mean values*** | ***Mean Diff.*** | ***95,00% CI of diff,*** | ***Significance*** | ***Adjusted P Value*** |
| 0 | 0 | 0 | 0 | -3,717 to 3,717 | ns | >0,9999 |
| 10 | 10.07 | 8.036 | 2.031 | -1,686 to 5,748 | ns | 0.8828 |
| 20 | 15.67 | 10.93 | 4.738 | 1,022 to 8,455 | ** | 0.0024 |
| 30 | 18.87 | 10.36 | 8.51 | 4,793 to 12,23 | **** | <0,0001 |
| 40 | 20.6 | 9.071 | 11.53 | 7,812 to 15,25 | **** | <0,0001 |
| 50 | 18.8 | 7.393 | 11.41 | 7,691 to 15,12 | **** | <0,0001 |
| 60 | 16.93 | 5.571 | 11.36 | 7,645 to 15,08 | **** | <0,0001 |
| 70 | 14.33 | 4.75 | 9.583 | 5,867 to 13,3 | **** | <0,0001 |
| 80 | 11.47 | 4.107 | 7.36 | 3,643 to 11,08 | **** | <0,0001 |
| 90 | 10 | 3 | 7 | 3,283 to 10,72 | **** | <0,0001 |
| 100 | 7.067 | 2.643 | 4.424 | 0,7072 to 8,14 | ** | 0.0066 |
| 110 | 4.8 | 2.429 | 2.371 | -1,345 to 6,088 | ns | 0.6796 |
| 120 | 4.6 | 2.036 | 2.564 | -1,152 to 6,281 | ns | 0.5396 |
| 130 | 3.4 | 2.214 | 1.186 | -2,531 to 4,902 | ns | 0.9998 |
| 140 | 3.067 | 2.071 | 0.9952 | -2,721 to 4,712 | ns | >0,9999 |
| 150 | 2.6 | 2.107 | 0.4929 | -3,224 to 4,209 | ns | >0,9999 |
| 160 | 2.467 | 1.821 | 0.6452 | -3,071 to 4,362 | ns | >0,9999 |
| 170 | 2 | 1.929 | 0.07143 | -3,645 to 3,788 | ns | >0,9999 |
| 180 | 1.933 | 1.75 | 0.1833 | -3,533 to 3,9 | ns | >0,9999 |
| 190 | 1.933 | 1.857 | 0.07619 | -3,64 to 3,793 | ns | >0,9999 |
| 200 | 1.933 | 1.714 | 0.219 | -3,498 to 3,936 | ns | >0,9999 |

| ***Sholl analysis statistics: 0-660 mm*** | | | | | | |
|---|---|---|---|---|---|---|
| | ***Ctrl mean values*** | ***Nr2f1 cKO mean values*** | ***Mean Diff.*** | ***95,00% CI of diff*** | ***Statistics*** | ***Adjusted P Value*** |
| 0 | 0 | 0 | 0 | -2,634 to 2,634 | ns | >0,9999 |
| 10 | 10.07 | 8.036 | 2.031 | -0,6032 to 4,665 | ns | 0.4705 |
| 20 | 15.67 | 10.93 | 4.738 | 2,104 to 7,372 | **** | <0,0001 |
| 30 | 18.87 | 10.36 | 8.51 | 5,875 to 11,14 | **** | <0,0001 |
| 40 | 20.6 | 9.071 | 11.53 | 8,894 to 14,16 | **** | <0,0001 |
| 50 | 18.8 | 7.393 | 11.41 | 8,773 to 14,04 | **** | <0,0001 |
| 60 | 16.93 | 5.571 | 11.36 | 8,728 to 14 | **** | <0,0001 |
| 70 | 14.33 | 4.75 | 9.583 | 6,949 to 12,22 | **** | <0,0001 |
| 80 | 11.47 | 4.107 | 7.36 | 4,725 to 9,994 | **** | <0,0001 |
| 90 | 10 | 3 | 7 | 4,366 to 9,634 | **** | <0,0001 |
| 100 | 7.067 | 2.643 | 4.424 | 1,79 to 7,058 | **** | <0,0001 |
| 110 | 4.8 | 2.429 | 2.371 | -0,2627 to 5,006 | ns | 0.1513 |

| | | | | | | |
|---|---|---|---|---|---|---|
| 120 | 4.6 | 2.036 | 2.564 | -0,06988 to 5,198 | ns | 0.0682 |
| 130 | 3.4 | 2.214 | 1.186 | -1,448 to 3,82 | ns | >0,9999 |
| 140 | 3.067 | 2.071 | 0.9952 | -1,639 to 3,629 | ns | >0,9999 |
| 150 | 2.6 | 2.107 | 0.4929 | -2,141 to 3,127 | ns | >0,9999 |
| 160 | 2.467 | 1.821 | 0.6452 | -1,989 to 3,279 | ns | >0,9999 |
| 170 | 2 | 1.929 | 0.07143 | -2,563 to 2,706 | ns | >0,9999 |
| 180 | 1.933 | 1.75 | 0.1833 | -2,451 to 2,818 | ns | >0,9999 |
| 190 | 1.933 | 1.857 | 0.07619 | -2,558 to 2,71 | ns | >0,9999 |
| 200 | 1.933 | 1.714 | 0.219 | -2,415 to 2,853 | ns | >0,9999 |
| 210 | 1.867 | 1.821 | 0.04524 | -2,589 to 2,679 | ns | >0,9999 |
| 220 | 2 | 1.929 | 0.07143 | -2,563 to 2,706 | ns | >0,9999 |
| 230 | 1.8 | 1.75 | 0.05 | -2,584 to 2,684 | ns | >0,9999 |
| 240 | 1.733 | 1.786 | -0.05238 | -2,687 to 2,582 | ns | >0,9999 |
| 250 | 1.8 | 1.964 | -0.1643 | -2,798 to 2,47 | ns | >0,9999 |
| 260 | 1.6 | 2 | -0.4 | -3,034 to 2,234 | ns | >0,9999 |
| 270 | 1.4 | 1.714 | -0.3143 | -2,948 to 2,32 | ns | >0,9999 |
| 280 | 1.333 | 1.607 | -0.2738 | -2,908 to 2,36 | ns | >0,9999 |
| 290 | 1.067 | 1.607 | -0.5405 | -3,175 to 2,094 | ns | >0,9999 |
| 300 | 1.133 | 1.786 | -0.6524 | -3,287 to 1,982 | ns | >0,9999 |
| 310 | 1.2 | 1.607 | -0.4071 | -3,041 to 2,227 | ns | >0,9999 |
| 320 | 1.333 | 1.25 | 0.08333 | -2,551 to 2,718 | ns | >0,9999 |
| 330 | 1.333 | 1.179 | 0.1548 | -2,479 to 2,789 | ns | >0,9999 |
| 340 | 1.333 | 1.25 | 0.08333 | -2,551 to 2,718 | ns | >0,9999 |
| 350 | 1.4 | 1.25 | 0.15 | -2,484 to 2,784 | ns | >0,9999 |
| 360 | 1.333 | 1.143 | 0.1905 | -2,444 to 2,825 | ns | >0,9999 |
| 370 | 1.333 | 1.107 | 0.2262 | -2,408 to 2,86 | ns | >0,9999 |
| 380 | 1.267 | 0.9643 | 0.3024 | -2,332 to 2,937 | ns | >0,9999 |
| 390 | 1.133 | 1.071 | 0.0619 | -2,572 to 2,696 | ns | >0,9999 |
| 400 | 1.067 | 1 | 0.06667 | -2,568 to 2,701 | ns | >0,9999 |
| 410 | 0.8667 | 0.9286 | -0.0619 | -2,696 to 2,572 | ns | >0,9999 |
| 420 | 0.8667 | 0.8571 | 0.009524 | -2,625 to 2,644 | ns | >0,9999 |
| 430 | 1.067 | 0.6071 | 0.4595 | -2,175 to 3,094 | ns | >0,9999 |
| 440 | 1.467 | 0.7143 | 0.7524 | -1,882 to 3,387 | ns | >0,9999 |
| 450 | 1.267 | 0.6786 | 0.5881 | -2,046 to 3,222 | ns | >0,9999 |
| 460 | 1 | 0.4286 | 0.5714 | -2,063 to 3,206 | ns | >0,9999 |
| 470 | 0.9333 | 0.25 | 0.6833 | -1,951 to 3,318 | ns | >0,9999 |
| 480 | 0.9333 | 0.1071 | 0.8262 | -1,808 to 3,46 | ns | >0,9999 |
| 490 | 0.8667 | 0.07143 | 0.7952 | -1,839 to 3,429 | ns | >0,9999 |
| 500 | 0.8 | 0.07143 | 0.7286 | -1,906 to 3,363 | ns | >0,9999 |
| 510 | 0.8 | 0.07143 | 0.7286 | -1,906 to 3,363 | ns | >0,9999 |
| 520 | 1.267 | 0.07143 | 1.195 | -1,439 to 3,829 | ns | 0.9999 |
| 530 | 0.9333 | 0.07143 | 0.8619 | -1,772 to 3,496 | ns | >0,9999 |
| 540 | 0.4667 | 0.07143 | 0.3952 | -2,239 to 3,029 | ns | >0,9999 |
| 550 | 0.2 | 0.07143 | 0.1286 | -2,506 to 2,763 | ns | >0,9999 |
| 560 | 0.1333 | 0.07143 | 0.0619 | -2,572 to 2,696 | ns | >0,9999 |

| | | | | | | |
|---|---|---|---|---|---|---|
| 570 | 0 | 0.07143 | -0.07143 | -2,706 to 2,563 | ns | >0,9999 |
| 580 | 0 | 0.07143 | -0.07143 | -2,706 to 2,563 | ns | >0,9999 |
| 590 | 0 | 0.07143 | -0.07143 | -2,706 to 2,563 | ns | >0,9999 |
| 600 | 0 | 0.07143 | -0.07143 | -2,706 to 2,563 | ns | >0,9999 |
| 610 | 0 | 0.07143 | -0.07143 | -2,706 to 2,563 | ns | >0,9999 |
| 620 | 0 | 0.07143 | -0.07143 | -2,706 to 2,563 | ns | >0,9999 |
| 630 | 0 | 0.07143 | -0.07143 | -2,706 to 2,563 | ns | >0,9999 |
| 640 | 0 | 0.03571 | -0.03571 | -2,67 to 2,598 | ns | >0,9999 |
| 650 | 0 | 0.03571 | -0.03571 | -2,67 to 2,598 | ns | >0,9999 |
| 660 | 0 | 0 | 0 | -2,634 to 2,634 | ns | >0,9999 |

***Table S2.*** *Analysis of basal dendritic complexity of layer V pyramidal neurons (LVPNs). Sholl analysis data of layer V pyramidal neurons in Ctrl and Nr2f1cKO brain slices of proximal (0-120mm) and total (0-660mm) arborization. P-values are calculated by 2way ANOVA test. See also Figure 4 and Figure S3.*

| Antigen | Provider | Catalog # | Species | Application | Working dilution |
|---|---|---|---|---|---|
| **AnkG** | Antibodies Inc. | 75-146 | Ms | IF - 1AB | 1:200 |
| **CTIP2** | Abcam | ab18465 | Rat | IF - 1AB | 1:500 |
| **NeuN** | Cell Signaling | 24307S | Rb | IF - 1AB | 1:100 |
| **HCN1** | Abcam | ab229340 | Rb | IF - 1AB<br>WB | 1:100<br>1:1000 |
| **b-actin** | Abcam | ab8277 | Rb | WB | 1:3000 |
| **GFP** | Abcam | ab13970 | Ck | ChIP | 3 mg |
| **COUP-TFI** | Thermo Fisher | PA5-21611 | Rb | ChIP | 3 mg |
| **Ms IgG - AF 555** | Thermo Fisher | A21422 | Gt | IF - 2AB | 1:500 |
| **Rat IgG - AF 488** | Thermo Fisher | | | IF - 2AB | 1:500 |
| **Rb IgG - AF 647** | Thermo Fisher | A32733 | Gt | IF - 2AB | 1:500 |
| **Rb IgG - AF 555** | Thermo Fisher | A21428 | Gt | IF - 2AB | 1:100 |
| **Ck IgY - AF 488** | Thermo Fisher | A11039 | Gt | IF - 2AB | 1:500 |
| **Rb IgG - HRP** | BioRad | 1706515 | Gt | WB - 2AB | 1:3000 |
| **Ms IgG - HRP** | BioRad | 1721011 | Gt | WB - 2AB | 1:3000 |

***Table S3***. *Primary and secondary antibody used in this study.*
